# Analysis of cell-type-specific chromatin modifications and gene expression in *Drosophila* neurons that direct reproductive behavior

**DOI:** 10.1101/2020.11.16.384461

**Authors:** Colleen M Palmateer, Shawn C Moseley, Surjyendu Ray, Savannah G Brovero, Michelle N Arbeitman

## Abstract

Examining the role of chromatin modifications and gene expression in neurons is critical for understanding how the potential for behaviors are established and maintained. We investigate this question by examining *Drosophila melanogaster fru P1* neurons that underlie reproductive behaviors in both sexes. We developed a method to purify cell-type-specific chromatin (Chromatag), using a tagged histone H2B variant that is expressed using the versatile Gal4/UAS gene expression system. Here, we use Chromatag to evaluate five chromatin modifications, at three life stages in both sexes. We find substantial changes in chromatin modification profiles across development and fewer differences between males and females. We generated cell-type-specific RNA-seq data sets, using translating ribosome affinity purification (TRAP), and identify actively translated genes in *fru P1* neurons, revealing novel stage- and sex-differences in gene expression. We compare chromatin modifications to the gene expression data and find patterns of chromatin modifications associated with gene expression. An examination of the genic features where chromatin modifications resides shows certain chromatin modifications are maintained in the same genes across development, whereas others are more dynamic, which may point to modifications important for cell fate determination in neurons. Using a computational analysis to identify super-enhancer-containing genes we discovered differences across development, and between the sexes that are cell-type-specific. A set of super-enhancer-containing genes that overlapped with those determined to be expressed with the TRAP approach were validated as expressed in *fru P1* neurons.

**Author Summary:** Differences in male and female reproductive behaviors are pervasive in nature and important for species propagation. Studies of sex differences in the fruit fly, *Drosophila melanogaster*, have been ongoing since the early 1900s, with many of the critical molecular and neural circuit determinates that create sexually dimorphic behavior identified. This system is a powerful model to understand fundamental principles about the underpinnings of complex behavior at high resolution. In this study, we examine the gene expression and chromatin modification differences specifically in a set of neurons that direct male and female reproductive behaviors in *Drosophila*. We describe differences across development and between the sexes with the goal of understanding how the potential for behavior is created and maintained.

## Introduction

Innate reproductive behaviors are directed by neurons that attain their cell fate during development. Cell fate identity is then maintained during adult stages for the performance of behavior. For neurons, plasticity and modification by experience are an intrinsic part of their function. Chromatin and DNA modifications have been implicated as a mechanism that could direct both long-lasting cell fate identity and allow for plasticity, by directing cell-type-specific transcriptional programs that can be modified in response to the environment. Indeed, studies have found chromatin and DNA modifications play a role in synaptic plasticity, learning, memory, stress response, and complex social behaviors across phyla (reviewed in 1, 2–5). Furthermore, DNA and chromatin modifications can provide a mechanism to direct stage-specific and sex-differences in behavior, by overlaying time- and sex-specific information onto the genome that is less transient than transcriptional changes. Here, we examine neurons underlying *Drosophila melanogaster* reproductive behaviors to investigate the dynamic developmental and sex-specific roles of chromatin modifications for altering gene expression in neurons that direct complex, innate behaviors.

Reproductive behaviors in *Drosophila* include a multi-step, male courtship display towards the female (6, reviewed in 7). If the female chooses to mate, she will slow down to allow mating to occur and subsequently will display post-mating behaviors that include egg-laying, and additional post-mating changes (reviewed in 8, 9). The potential for these sexually dimorphic behaviors is specified by the sex hierarchy transcription factors encoded by *doublesex* (*dsx*) and *fruitless* (*fru*) (10–15) (**Figure 1A**). Sex-specific Dsx and Fru are produced in subsets of neurons important for reproductive behaviors (< 5% of neurons). These transcription factors are first detected in the nervous system at late third instar larvae when the nervous system begins to be remodeled for adult function, are present throughout metamorphosis, and persist into adult stages (14–20).

**Figure 1.**
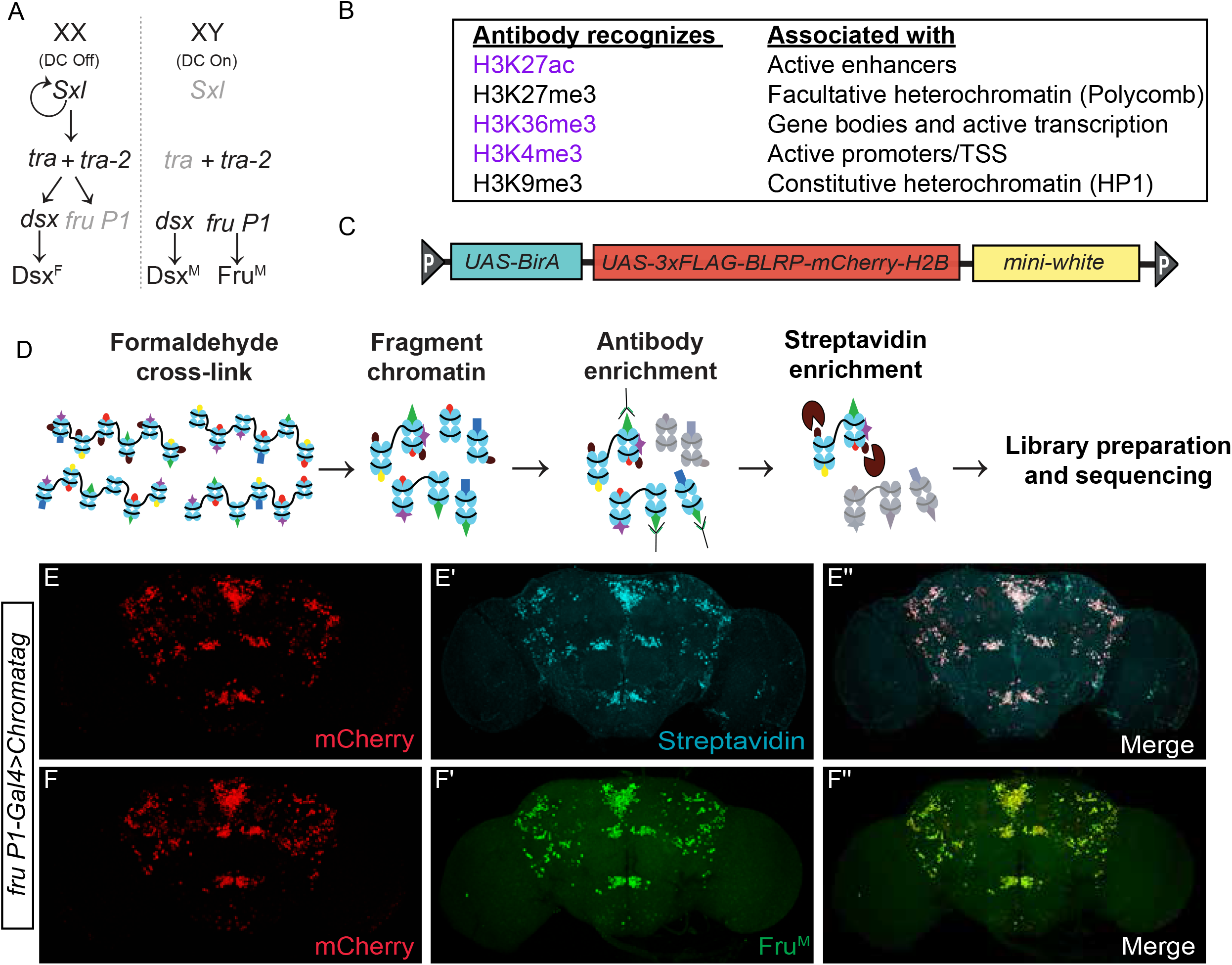
Chromatag methodology enables cell-type-specific ChIP in *fru P1* neurons. **(A)** The *Drosophila* somatic sex determination hierarchy is composed of an alternative pre-mRNA splicing cascade dependent on the number of X chromosomes. Alternative pre-mRNA splicing of *doublesex* (*dsx*) and *fruitless* (*fru P1*) products generates sex-specific transcription factors: Dsx^F^, Dsx^M^, and Fru^M^. Sex-specific Dsx and Fru regulate gene expression to yield sex differences in morphology, neuroanatomy and ultimately behavior (6, 7, reviewed in 82) **(B)** Histone H3 modifications profiled in this study and their role in transcriptional activation (purple) or repression (black) (reviewed in 36, 37–40). **(C)** Schematic of the *Chromatag* transgene. The construct includes three transgenes. One directs expression of amino-terminal tagged H2B under UAS control; the tag contains 3x-FLAG, the <u>B</u>iotin <u>L</u>igase <u>R</u>ecognition <u>P</u>eptide (BLRP) and *mCherry*. Another transgene directs biotin ligase (*BirA*) expression under UAS control. The *mini-white* transgene is a germline transformation marker. **(D)** Overview of the Chromotag-ChIP workflow. Tissue from flies expressing Chromatag is fixed with formaldehyde, the chromatin is isolated and then fragmented by sonication. An antibody to a histone modification is used in the first ChIP. A subsequent pull-down with streptavidin-bound beads enriches for cell-type-specific chromatin. The purified genomic DNA undergoes library preparation and sequencing. **(E-F”)** Immunostaining of 10-12 day adult male brains expressing the *Chromatag* transgene in *fru P1* neurons. **(E-E”)** The Chromatag H2B variant is visualized by the mCherry tag (red) and overlaps with biotinylated H2B detected using fluorescently conjugated streptavidin (cyan). **(F-F’’)** The Chromatag H2B variant is visualized by the mCherry tag (red) and overlaps with Fru^M^ based on anti-Fru^M^ immunostaining (green). **(E-F”)** Flies are the genotype: *w/Y; P[w*^+*mC*^, *UAS-GAL4]/P[w*^+*mC*^, *UAS-Chromatag]; fru P1-Gal4*/+.

*fru* was first identified as important for reproductive behavior based on mutants that had altered male courtship behaviors (21). In-depth, molecular-genetic analysis showed that *fru* is a complex locus containing at least four promoters (P1-P4), with the *fru P1* promoter driving expression of a transcript class (*fru P1*) that is spliced downstream of the sex determination hierarchy (**Figure 1A)** (11, 12). Sex-specific splicing of *fru P1* transcripts results in production of male-specific transcription factor isoforms (Fru^M^) that are members of the BTB family, whereas *fru P1* transcripts in females have an early stop codon, with a coding potential of 94 amino acids of unknown function (11, 12). Fru^M^ isoforms contain an additional amino-terminal 101 amino acid region with no known function, differentiating them from common Fru isoforms produced by P2-P4 promoters (11, 12). Mutations impacting *fru P1* transcripts alter several aspects of male courtship, but do not appear to alter female behaviors, though some allele combinations showed female sterility (11, 12, 22–24). Additionally, forcing male-specific splicing or producing Fru^M^ in females allows for some male courtship behaviors in females (19, 25). However, even though *fru P1* products have a major role in only male behavior, *fru P1*-expressing neurons (*fru P1* neurons hereafter) are present in both sexes and direct the distinct male and female reproductive behaviors (19, 20, 25, 26). A major remaining question is how does a largely shared *fru P1* neuroanatomical substrate and shared genome give rise to different male and female reproductive behaviors.

It is clear that *fru P1* neurons and *fru P1* expression are modified during both developmental and adult stages. Sex differences in the number and morphology of some *fru P1* neurons has been shown to be established during metamorphosis (17, 20, 27). Furthermore, early genetic studies using a temperature sensitive allele of the sex hierarchy gene *transformer2* that regulates splicing of *fru P1* showed that the middle of metamorphosis was a critical period for establishing sex-specific behaviors, and also showed that there was adult plasticity in terms of behavioral potential (28). Later studies, performed by overexpressing the female-splice-form of the sex hierarchy gene *transformer* (*tra*), which also regulates splicing of *fru P1,* showed that there is an irreversible period critical for establishing the potential for sex-specific behavior during early metamorphosis (29). In the adult, *fru P1* expression is regulated in *Or47b* and *Ir84a* olfactory receptor neurons (ORNs) and is dependent on adult olfactory activity (30). Additionally, an increase in adult *fru P1* expression in *Or47b* ORNs has been linked to an increase in activating chromatin modifications at the *fru P1* promotor (31). Further, *fru P1* expression in adult *Or47b* ORNs is required for increased copulation rate in group housed males compared to single housed males (32). These studies point to developmental and adult stages as critical for establishing the potential for behavior. Therefore, an examination of the molecular-genetic basis of reproductive behavior directed by *fru P1* should include understanding of both developmental and adult stages.

One mechanism to generate and maintain behavioral potential is through chromatin modifications. A previous study demonstrated that Fru^M^ forms a complex with transcriptional cofactor, Bonus, a TIF1 homolog (33). This Fru^M^-Bonus complex selectively recruits the chromatin modifying protein Histone deacetylase 1 (HDAC1/Rpd3) or Heterochromatin protein 1a (HP1a/Su(var)205) (33). If the Fru^M^-Bonus complex recruits HDAC1, a set of sexually dimorphic interneurons (mAL) are masculinized, whereas recruitment of HP1a results in demasculinized morphology of the mAL neurons (33). This suggests that differences in chromatin modifications in *fru P1* neurons may contribute to gene expression changes. Here, we ask about differences in the chromatin landscape in male and female *fru P1* neurons, how chromatin modifications change across developmental and adult stages, and if chromatin modifications are a mechanism to direct stage- and sex-specific gene expression that direct behavior.

To determine the chromatin landscape in *fru P1* neurons, we developed a transgenic strategy to purify chromatin using a tagged histone H2B, under Gal4/UAS transcriptional control (called Chromatag). Complementary to other cell-type-specific chromatin purification methods, Chromatag allows for the direct purification of chromatin and profiling of chromatin modifications without requiring FACS purification (reviewed in 34, 35). Using this method, we purified chromatin from *fru P1* neurons of both sexes and examine five chromatin modifications (H3K27ac, H3K36me3, H3K4me3, H3K27me3, and H3K9me3), with conserved roles in regulating gene expression (**Figure 1B**) (reviewed in 36, 37–40).We examine chromatin modifications at three stages: mid-metamorphosis at 48 hours after puparium formation (48hr APF), 1-day adults (frozen 16-24hrs post-eclosion), and 10-12-day adults, to gain insight into how the potential for behavior is established and maintained. In addition, we examined chromatin modifications in all neurons within the head (*elav*-*Gal4*) in 1-day adults to compare with neurons from 1-day adult *fru P1* neurons. To assess if different combinations of H3 modifications are correlated with transcriptional activity in *fru P1* neurons, we generate and perform a meta-analysis with data from cell-type-specific transcriptional studies (<u>t</u>ranslating <u>r</u>ibosome <u>a</u>ffinity <u>p</u>urification technique; TRAP) (41, 42). We observe greater changes in chromatin modification profiles across developmental stages than between sexes. We find that a subset of histone modification patterns is characteristic of expressed genes but that no pattern is indicative of high expression. Using our H3K27ac modification peaks, we have identified super-enhancer (SE) regions and show that they are largely stage specific. Informed by SE-containing genes and genes identified in our *fru P1* TRAP data sets, we selected genes and examined their expression in *fru P1* neurons. Altogether, we discovered critical chromatin modifications and gene expression patterns that are time point and sex-specific that underlie the potential for complex behavior.

## Results

### Cell-type-specific strategy to analyze chromatin modifications

To examine the chromatin modification landscape in a cell-type-specific manner we developed transgenic strains to express tagged histone H2B under Gal4/UAS control (43). This approach is versatile and can be used with other Gal4 drivers (reviewed in 44) to purify chromatin from different cell-types. The transgene contains a cDNA, under UAS control, for expression of H2B tagged with amino-terminal 3xFLAG, biotin ligase recognition peptide (BLRP), and mCherry (Chromatag; **Figure 1C**, red). The transgene also contains the *E. coli* biotin ligase gene, *birA*, under UAS control (45) (**Figure 1C**, blue). To perform cell-type-specific ChIP-seq, a sequential, two-step chromatin immunoprecipitation procedure was used, where first we enriched for chromatin fragments containing a histone modification, using an antibody purification, and then enriched for chromatin fragments from the cell-type of interest, using streptavidin purification (**Figure 1D**).

We first validated that the transgene is expressed in a cell-type-specific manner, by examining *UAS*-*Chromatag* expression driven by *fru P1-Gal4* (*fru P1-Gal4*>*Chromatag*). We show that mCherry-tagged H2B overlaps with Fru^M^ immunostaining, and that fluor-conjugated streptavidin staining is restricted to cells where mCherry-H2B is detected (**Figure 1E-F**), at all time points examined (**Supplemental Figure 1**). To confirm that Chromatag can be incorporated throughout the genome, we performed sequential ChIP-seq with an antibody to histone H3. As expected, we observed high coverage across all chromosomes (78-91% coverage), at all time points and in both cell-types examined (**Supplemental Table 1**). It is not possible to uniquely align short-read ChIP-seq data to repetitive sequence regions, which accounts for not having closer to 100% coverage. For example, we observed the lowest coverage on the 4^th^ chromosome which has highly repetitive DNA sequence content (reviewed in 46, reviewed in 47).

To determine if there are behavioral impacts of producing Chromatag in neurons, we examined the locomotor activity of *fru P1-Gal4*>*Chromatag* flies. There are no significant differences in locomotor activity between single transgene control flies and *fru P1-Gal4*>*Chromatag* flies (**Supplemental Figure 2A**). Additionally, we assayed courtship behavior in *fru P1-Gal4>Chromatag* flies. We find that *fru P1-Gal4*>*Chromatag* males court females normally and are similar to single transgene controls. We do find elevated male-male courtship in *fru P1-Gal4*>*Chromatag* flies, compared to single transgene controls, which is often seen with transgenic strains harboring the mini *white* cDNA, used as a marker for germline transformation (48–51).

### Correlation across chromatin modification data sets

In this study we generated ChIP-seq data sets for five histone H3 modifications in *fru P1* and *elav* neurons from both sexes (**Figure 1B**, **Supplemental Figure 1A**). We examined three activating modifications (H3K27ac, H3K4me3 and H3K36me3), and two repressive modifications (H3K27me3 and H3K9me3), with 3-4 biological replicates for each stage and sex. Using the deepTools package, ChIP-seq read count signals for each data set were normalized to the matched ChIP-seq input libraries in 50bp bins and the log2 ratio of signal was calculated (52). We next used the Spearman correlation heatmap function in deepTools to evaluate the positional correlation of genome-wide ChIP-seq signals from all the data sets comparing 10kb bins (**Supplemental Figure 3**), with color indicating correlation. The results show that the data sets for the same H3 histone modification are nearly all positively correlated (**Supplemental Figure 3**, blue regions in heatmap). The datasets for the two repressive modifications (H3K9me3 and H3K27me3) cluster together (pink highlighted node), as might be expected given these modifications are known to decorate heterochromatic regions (reviewed in 36, 37). The strongest anti-correlation occurs between the samples from adult stages for H3K27ac and H3K36me3 (light blue and dark blue nodes in dendrogram), indicating that these modifications are occurring in different gene regions and/or genes. The anti-correlation of H3K27ac and H3K36me3 is also consistent with H3K27ac residing in promotor regions and H3K36me3 in gene bodies (53, reviwed in 54). Additionally, these results show that the Chromatag approach is effective in cell-type-specific chromatin enrichment, given that the male and female data sets for each cell-type (*elav* and *fru P1*) are highly correlated with each other, in a stage-specific manner. Furthermore, the correlation analysis provides validation, as data for the same chromatin modification largely cluster together and show the cell-type specificity of the Chromatag approach. This analysis indicates that the distribution for each chromatin modification shows time point, cell-type and sex differences.

### Examination of chromatin modification peaks reveal differences in genomic distributions and genes in cell-types and across time points

Next, we used the MACS2 peak calling algorithm to identify genome regions enriched for each chromatin modification, using pooled biological replicate data (**Supplemental Table 1**) (55). We limited our analysis to peaks that are within three kilobases (kb) of an annotated gene, resulting in 16,385 genes containing at least one peak, across the five modifications (**Supplemental Table 1**). We note that even though we are able to effectively ChIP and sequence DNA from the male X-chromosome, the MACS2 peak calling algorithm called very few peaks for the male X-chromosome (**Supplemental Table 1**). To attempt to correct this inability to call peaks on the male X-chromosome, we adjusted several MACS2 model parameters and settings (see methods), and tried additional peak calling algorithms, but were not able to identify many additional peaks on the male X-chromosome (data not shown). We speculate that this may be because of the overall chromatin decondensation and conformational differences of the male X-chromosome due to dosage compensation, leading to difficulties in peak calling (56, reviwed in 57).

We determined where peaks occur, with respect to eleven gene features, and how the distribution of peaks changes proportionally over time (**Figure 2A-J**). We generally find that the proportion of peaks in the gene features are highly similar between males and females at each time point, for each histone modification, even though we cannot call many peaks on the male X-chromosome. This is true for the ChIP-seq data from both the *fru P1* and *elav* data sets. Given the limitation of calling peaks on the male X-chromosome, we next compared gene feature peak position within sex. In *fru P1* neurons, H3K27ac and H3K4me3 peaks show the largest differences across time points (**Figure 2A-J**). For example, the male and female data from 48hr APF have ~65% of H3K27ac peaks less than one kb from promotor regions, whereas in adult stages ~34% are less than one kb from the promoter, with a larger percentage of peaks now found in exons, introns, and intergenic regions (**Figure 2A-B**). Similarly, for H3K4me3 at 48hr APF in *fru P1* neurons, the majority of H3K4me3 peaks in both males (~95%) and females (~70%) are one kb from promotor regions. Then, the percentage of H3K4me3 peaks with positions distant from the promotor increases in 1-day adults and subsequently decreases at 10-12 days, with the changes more pronounced in females (**Figure 2G-H**). For H3K27me3, H3K36me3, and H3K9me3 we observe smaller changes in the proportion of peaks for the gene features across developmental time (**Figure 2C-F** **and** **I-J**).

**Figure 2.**
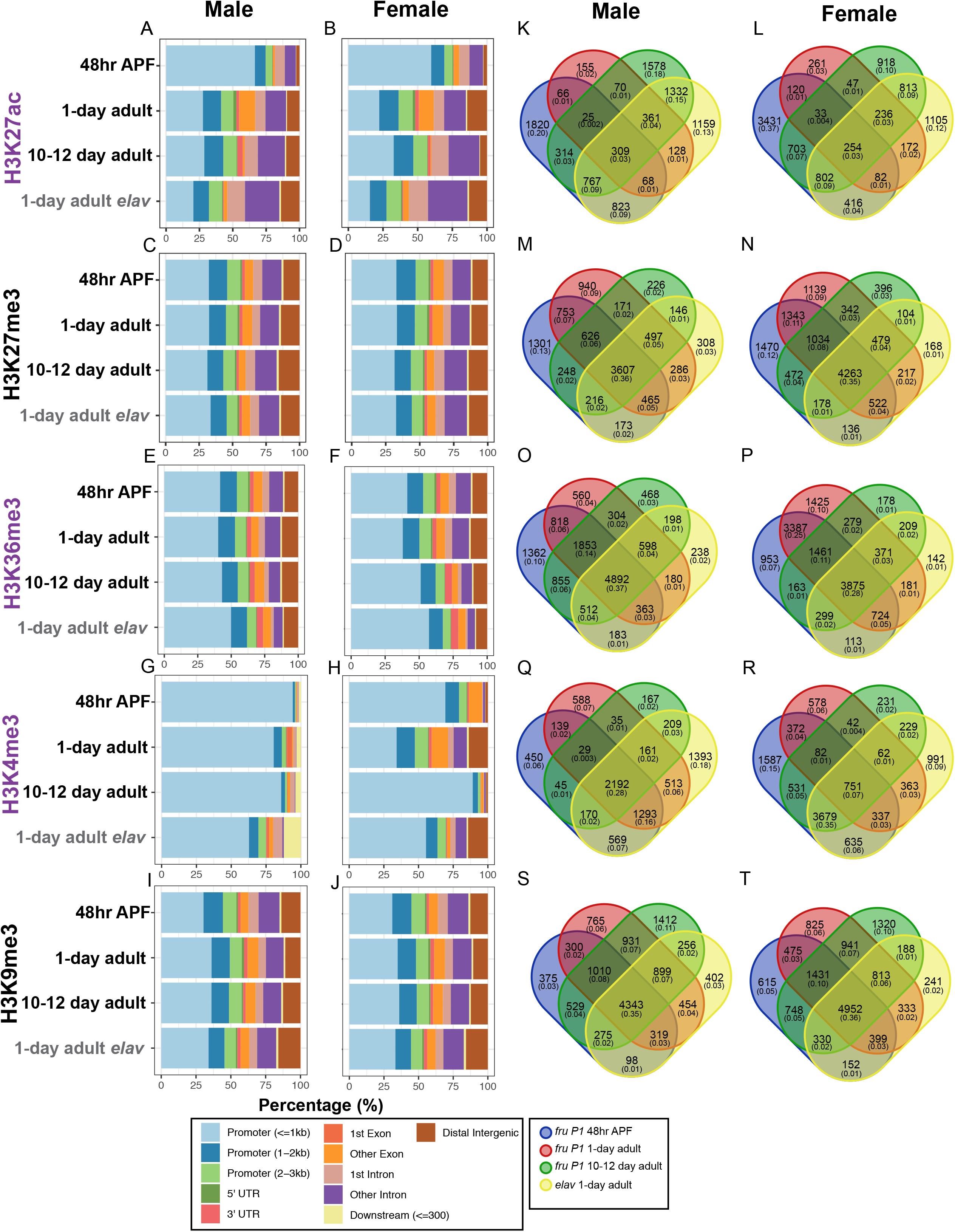
Genomic feature distribution and overlap of genes containing histone H3 chromatin modifications. **(A-J)** Genomic feature distributions of MACS2 peaks for activating (H3K27ac, H3K36me3, H3K4me3; purple) and repressive (H3K27me3 and H3K9me3; black) H3 chromatin modifications. The histone modification is indicated on the left. Each panel includes data for *fru P1* (black) and *elav* neurons (gray) in males (left) and females (right). The X-axis shows the percent of each genomic feature defined in the legend (bottom). **(K-T)** The histone modification is indicated on the left. Venn diagrams showing overlap of genes containing MACS2 peaks across time points in *fru P1* and *elav* neurons. The male (left) and female (right) data are indicated at top. The legend for the Venn diagrams is shown at bottom. For each Venn diagram category, the number of genes, and the proportion of the total genes (in parentheses) in each panel is shown. All MACS2 peaks were called on pooled biological replicates (n=3-4) and identified as enriched relative to matched input ChIP-seq libraries for each time point, sex, and neuron type. All MACS2 peaks in this analysis are listed in **Supplemental Table 1**.

Given that some modifications have substantial changes in the distribution of peaks over developmental time in *fru P1* neurons (H3K27ac and H3k4me3), we next determined if this is due to chromatin modifications in different genes or in the same genes, including both the *fru P1* and *elav* data sets in the analysis (**Figure 2K-T)** and in *fru P1* neurons only (**Supplemental Figure 4**). The observation that peaks are in different genes for the 1-day time point for the *fru P1* and *elav* chromatin data sets shows that there are cell-type-specific genes with each histone modification. We generated Venn diagrams to examine the overlap in *fru P1* neurons only (**Supplemental Figure 4**). Across the three time points, the genes with H3K27ac peaks have low overlap in males (4% overlap) and in females (3% overlap). The genes with H3K4me3 peaks have more overlap across the three time points in males (34% overlap), than females (9% overlap), which may be related to the observation that the distribution of the peaks have fewer changes in males over time (**Supplemental Figure 4**). For the modifications where the distribution of gene feature distribution does not change substantially over the three time points (H3K27me3, H3k36me3, and H3K9me3), there is also more overlap across the three time points for genes with these modifications (range is 40-51% overlap; **Supplemental Figure 4**). The H3K27ac and H3k4me3 activating modifications are largely found in different genes across time points and thus are not likely to be mediating cell fate determination, whereas the H3K27me3, H3k36me3, and H3K9me3 modifications persist across three time points, suggesting these chromatin modifications may be part of a mechanism to direct cell fate determination.

### Hierarchical clustering of genome-wide chromatin modifications in *fru P1* and *elav* neurons reveal stage- and sex-dependent differences

Next, using hierarchical clustering we examined the combinations of histone modifications on all annotated genes (**Figure 3 and Supplemental Table 2**). The heatmaps show clustered genes that are grouped based on similarity across the five chromatin modifications. This was performed using the input normalized data generated using deepTools, which calculates enriched and depleted ChIP regions over 50bp bins, relative to input, and is effective genome wide. The heatmaps show the ChIP enrichment and depletion, 3kb upstream of the start of transcription (TSS), the length of the normalized gene body (shown as 5kb) and 3kb downstream of the transcriptional end (TES).

**Figure 3.**
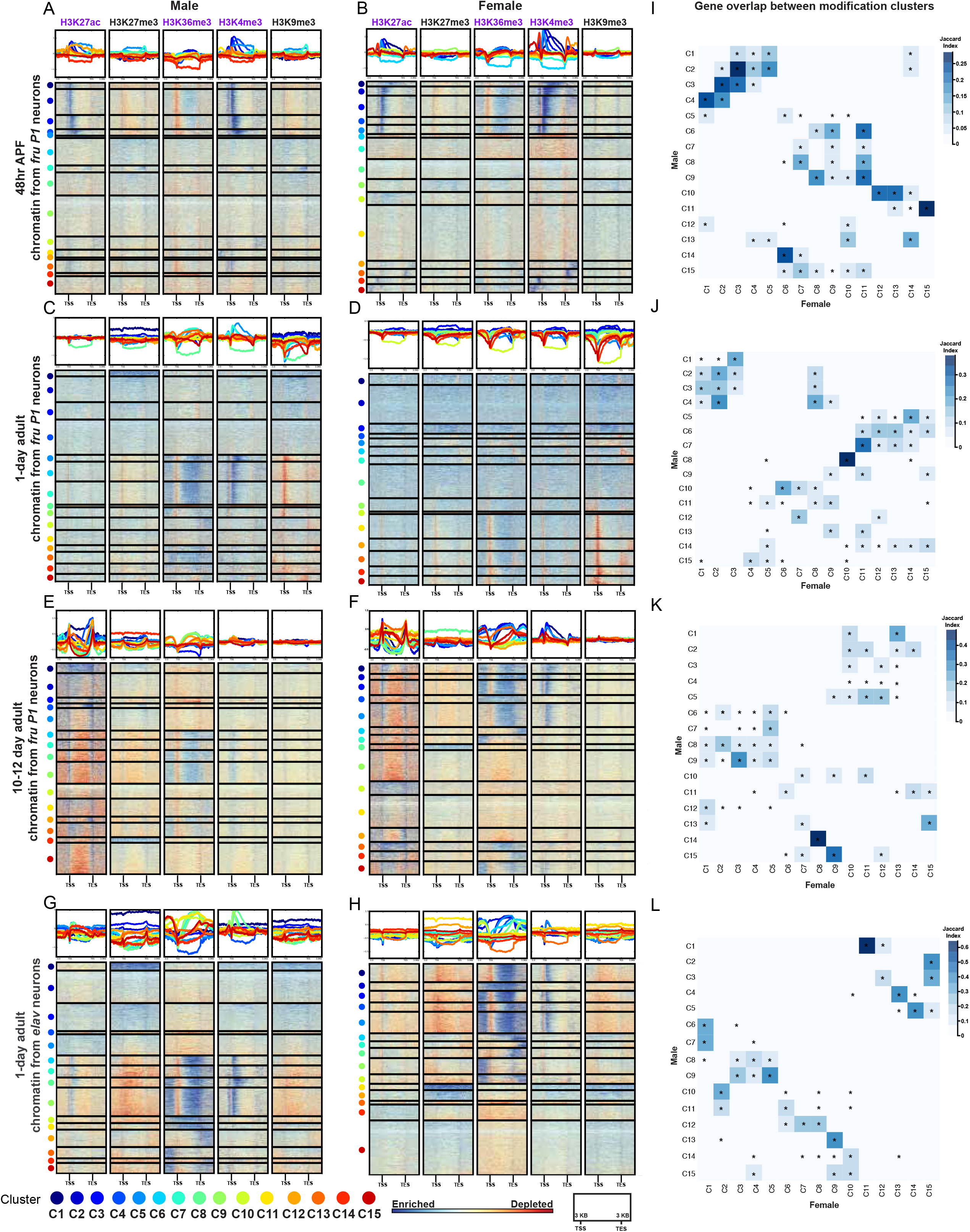
Genome-wide hierarchical clustering of histone modification distributions and cluster overlaps between the sexes. **(A-F)** Hierarchal clustering reveals shared combinations of histone modifications on all annotated genes, generated using deepTools. The ChIP enrichment of the input normalized data is indicated on the color bar, from blue (enrichment) to red (depletion), calculated in 50bp bins. The genes (rows) are clustered by the similarity of the ChIP patterns across all five histone modifications (each cluster number marked with a colored dot). The heatmap is divided into five columns, one for each chromatin modification. The top panel shows the average distribution of a chromatin modification in each cluster, with the line color indicating the cluster number. For each chromatin modification, the heatmap shows scaled gene bodies (scaled to 5kb) and 3kb upstream and downstream of the transcriptional start site (TSS) and transcriptional end site (TES). Heatmaps for *fru P1* chromatin data from 48hr APF **(A-B)**, 1-day **(C-D)**, and 10-12 day **(E-F)**. Heatmaps for *elav* chromatin data from 1-day **(G-H)**. The sex is indicated at the top. **(I-L)** GeneOverlap heatmap (61), where color indicates the Jaccard Index score, showing the similarity of the gene lists between clusters in males and females. For each panel the X-axis indicates female cluster number and the Y-axis indicates the male cluster number (color legend for clusters on the bottom). The legend for the Jaccard similarity index is on the right of each panel. Asterisks indicate statistically significant overlap of genes between clusters (Fisher’s exact test, p<0.05). Gene list information for all heatmap clusters is available in **Supplemental Table 2**. Gene overlap index and statistics are available in **Supplemental Table 6**.

First, these heatmaps provide further validation of the approach, given that overall the positions where there are enrichments of ChIP signal are consistent with the known locations for these modifications in *Drosophila* (53, 58). For the activating modifications, we find H3K27ac near the TSS, H3K4me3 near the TSS, and H3K36me3 along the gene body. Both repressive modifications show less localized enrichment patterns, which may be because we are examining genes, and not pericentric and repetitive regions which are typically heterochromatic (reviewed in 59). We do find that H3K9me3 has pronounced depletion at the TSS of some genes at the 1-day time point, which is unique compared to other stages examined here. One unexpected finding is the presence of H3K27ac peaks at the TES in 10-12 day adults, which to our knowledge is not widely reported.

Overall, the enrichment patterns of activating chromatin modifications in *fru P1* neurons show the largest differences across time points and not between the sexes, which is not unexpected given that most *Drosophila* genes do not have sex-specific expression patterns in *fru P1* neurons (see TRAP section below). For example, in both sexes at 48hr APF, H3K27ac enrichment mainly surrounds the TSS, in the sets of clusters with most pronounced ChIP-seq enrichment (**Figure 3A-B**, dark blue). At 1-day, the H3K27ac enrichment is more diffuse in pattern (**Figure 3C-D**). By the 10-12 day time point, H3K27ac is enriched near both the TSS and TES (dark blue), and this is observed for the majority of genes (**Figure 3 E-F**). In addition, in both sexes at 48hr APF, the genes with the most pronounced H3K27ac enrichment near the TSS (dark blue) also have the most pronounced H3K4me3 enrichment near the TSS and along the gene body (dark blue) (**Figure 3A-B**). Furthermore, H3K36me3 shows changes over developmental time, with high enrichment along the entire gene body seen at the later two time points and is not as pronounced at the 48hr APF time point (**Figure 3A-F**).

The repressive modifications also show time point specific differences. Notably, there is a set of genes with the H3K27me3 mark along the length of the gene found only in the adult data sets (**Figure 3C-H**; **C**-C1, **D**-C3, **E**-C14, **F**-C8, **G**-C1, and **H-**C11). Interestingly, gene ontology enrichment analysis to determine if there are overrepresented genes with known functions reveals that the list is enriched with “homeobox domain” encoding genes (60), which suggests that these critical transcriptional regulators are silenced after pupal stages in neurons, consistent with previous findings (58) (**Supplemental Tables 2 and 4**). Additionally, there are pronounced changes in the pattern of H3K9me3 over developmental time, with the 1-day time point having the most defined patterns of enrichment and depletion over the gene regions examined (**Figure 3 C-D**).

We also observe minor sex- and cell-type-specific differences in chromatin modifications, at the adult stages. At the 1-day time point, there is H3K4me3 enrichment at the TSS in males that is not as pronounced as in females (**Figure 3C-D**). Then at the 10-12 day time point, there is more pronounced H3K4me3 enrichment near the TSS in females (**Figure 3E-F**). The sex differences described are not observed in the *elav* neuron data, consistent with *fru P1* neurons having sex-specific identity (**Figure 3C-D** **and** **G-H**). The overall pattern between *fru P1* neurons and *elav* neurons at the 1-day time point are different, with all the modifications in the *elav* data showing more pronounced enrichment and depletion along the gene regions examined, for many genes (**Figure 3C-D** **and** **G-H**).

We next determined if the male and female clusters with similar patterns of histone modifications contain the same genes. To do this, for each time point we examined the overlap of genes between clusters from both sexes, using the GeneOverlap analysis tool (**Figure 3I-L**, **Supplemental Table 3**) (61). The blue color scale indicates the Jaccard similarity coefficient and the asterisk indicates the significance of the overlap based on a Fisher’s exact test (*p*<0.05). The clusters in the heatmap with the most overlapping genes between the sexes tend to have similar patterns of histone modification enrichment (**Figure 3I-L**, darkest blue boxes). For example, at 48hr APF, clusters 1-4 in males, and clusters 1-5 in females have similar H3K27ac, H3K36me3 and H3K4me3 patterns and several of these clusters have significant overlap in genes (**Figure 3A-B** **and** **I**). We also find clusters that have significant overlap between males and females but have different patterns of histone modifications (**Figure 3I-L**). One example of this is at 48hr APF, where there is a significant overlap of genes between male cluster 11 and female clusters 13 and 14, but different patterns of enrichment for H3K27ac and H3K4me3 (**Figure 3A-B**, **Figure I**). However, male cluster 11 also has significant gene overlap with female cluster 15 and the same patterns of histone modifications. *fru P1* chromatin data sets have more clusters with significant gene overlap between the sexes than the *elav* chromatin data sets (**Figure 3I-L**). In all adult *fru P1* and *elav* data sets, the clusters with repressive modifications containing the homeobox genes have a highly similar set of genes between males and females (**Figure 3 C-H** **and** **J-L**). Overall, we observe that the majority of clusters with a similar modification enrichment pattern in both sexes share a significant number of genes. There is a smaller subset of clusters with significant gene overlap that have different histone modification profiles.

### TRAP RNA-seq identifies mRNAs that are enriched in *fru P1* neurons in a stage-specific manner

Next, we examined gene expression specifically in *fru P1* neurons from males and females, using the translating ribosome affinity purification (TRAP) approach (41). The TRAP approach allows one to enrich for mRNAs that are actively translated in a cell-type-specific manner, via Gal4/UAS expression of a GFP-tagged ribosomal subunit (RpL10a). We examined all *fru P1* neurons from the 48hr APF stage and *fru P1* neurons from adult heads at the 10-12 adult stage, which is equivalent to the *fru P1* neurons profiled in the chromatin data sets. We have previously described *fru P1* TRAP gene expression results for neurons from 1-day adult heads and present those results in comparisons here (42). For each time point, we performed gene expression analysis at the exon-level, to account for transcript-isoform expression differences. A gene was considered differentially enriched, if at least one exon had differential expression (FDR <0.2).

At each time point, to identify genes with enriched expression in *fru P1* neurons, in either males or females, we performed two statistical comparisons: 1) male TRAP vs. male total mRNA; and 2) female TRAP vs. female total mRNA, which is similar to the comparisons for the chromatin data sets (hereafter called TRAP-enriched; **Supplemental Table 5**). We also present sex differences in either TRAP or total mRNA: 1) male TRAP vs. female TRAP (hereafter TRAP-biased); and 2) male total mRNA vs. female total mRNA (**Supplemental Table 6**). As expected, some *fru* exons, including the *fru P1*-specific exons, are enriched in the TRAP samples, relative to total mRNA control (**Supplemental Table 5**). In 48hr APF TRAP vs. total mRNA, the *fru P1* specific exons are enriched in TRAP samples, but do not meet the threshold of significance, whereas other *fru* exons show significant enrichment (**Supplemental Table 5**). In 10-day TRAP vs. total mRNA, *fru P1* and other *fru* exons are significantly enriched in TRAP samples, providing overall validation of the approach (**Supplemental Table 5**). When we examine the mapped reads at the *fru* locus at 48hr APF and 10-12 days, the sex-specifically spliced gene region shows the expected abundance of reads, with more in females, since the retained exonic region is larger in females (**Supplemental Figure 5**), similar to our observations at 1-day (42). Additionally, as further confirmation of the approach, the male and female TRAP-enriched and TRAP-biased gene lists significantly overlap with lists of genes that are predicted to be direct Fru^M^ targets (62, 63) (**Supplemental Table 3**). Here, an analysis of the genes that are more highly expressed in *fru P1* neurons vs. total mRNA (TRAP-enriched) and TRAP male and female-biased genes (TRAP-biased) is presented. The other gene lists from additional comparisons are provided (**Supplemental Tables 5 and 6**).

First, we compared the number of genes with enriched expression in *fru P1* neurons that are male TRAP-enriched, female TRAP-enriched, or TRAP-enriched in both sexes, using Venn diagrams (**Figure 4A** **and Supplemental Table 5**). Given that each time point has a different overall number of genes with TRAP-enriched expression, we also report the proportions of genes with enriched expression in each category to facilitate comparisons across the time points. We find a higher percent of enriched genes in male TRAP-enriched lists than female TRAP-enriched lists at 48hr APF and in 1-day adults (**Figure 4A**). At 48hr APF, 34% of genes are male TRAP-enriched, 12% are female TRAP-enriched and 54% are enriched in both male and female TRAP samples. At 1-day, 53% of genes are male-enriched,13% are female-enriched and 33% are in enriched in both male and female TRAP samples. In contrast, at the 10-12-day time point, a large fraction of genes have TRAP-enriched expression in both sexes (74%) and fewer male TRAP-enriched (6%) than female TRAP-enriched (20%). While there is a shared set of genes between males and females with *fru P1* TRAP-enriched expression at each stage, 48hr APF and 1-day adults show a higher fraction of genes with male TRAP-enriched expression. This is consistent with our previous observations where we suggest that this could be due to Fru^M^ gene regulation in males (42). This result is also in line with previous work showing *fru P1* has a critical function during development for reproductive behaviors (28, 29, 64).

**Figure 4.**
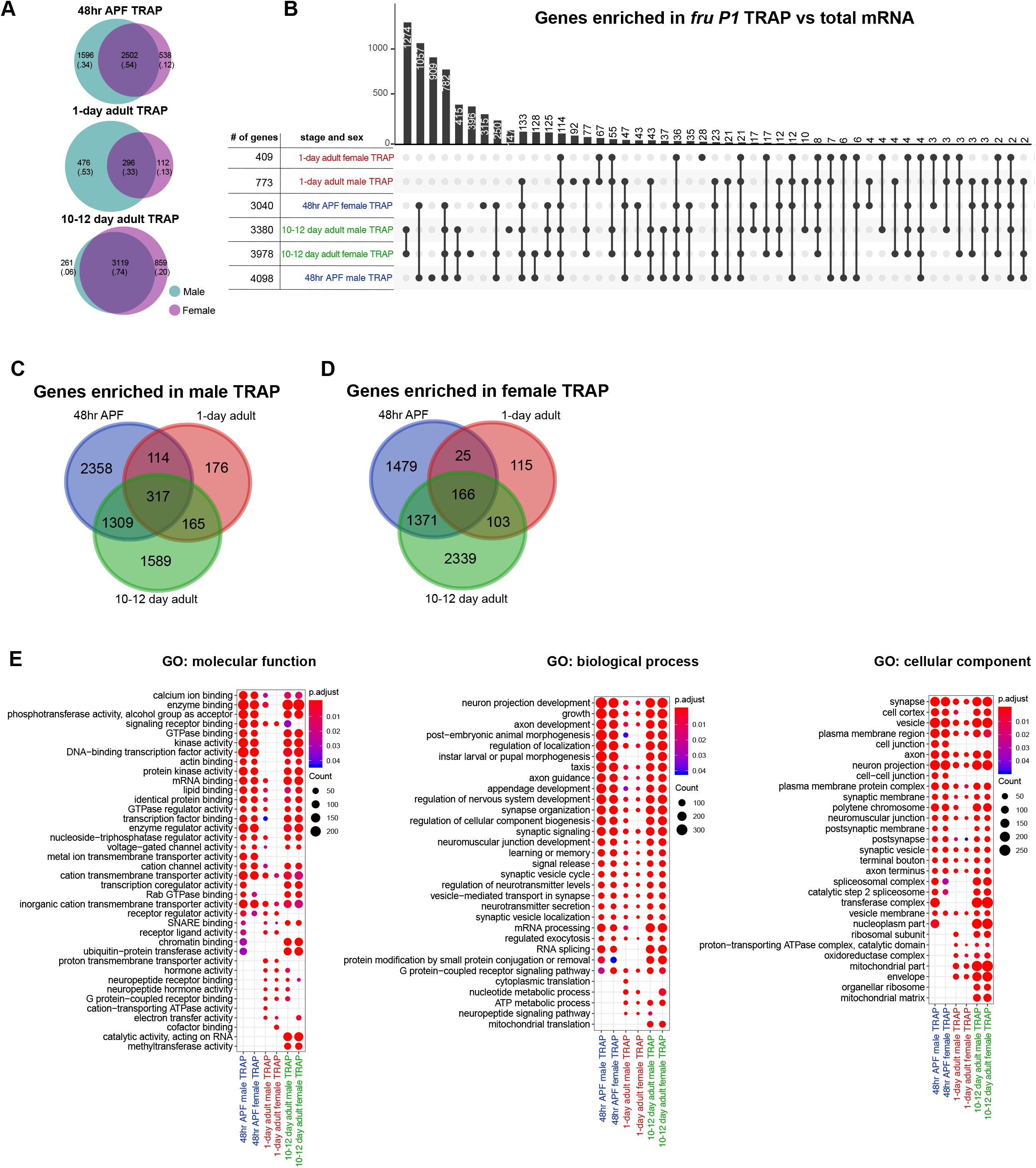
TRAP RNA-seq identifies mRNAs that are enriched in *fru P1* neurons at multiple time points. Analysis of TRAP-enriched genes. These genes were identified based on having at least one exon with significantly higher expression in TRAP vs. total mRNA (FDR<0.2). **(A)** Venn diagrams compare TRAP-enriched genes between males (teal) and females (purple) at three time points. For each Venn diagram category, the number of genes, and the proportion of the total genes (in parentheses) in each panel is shown. **(B)** Upset intersection plot of TRAP-enriched genes from 48hr APF (blue), 1-day (red) and 10-12 day (green) time points. n=4-5 biological replicates per condition. (**C-D**) Venn diagrams comparing TRAP-enriched genes across time points for each sex. (**E**) Gene ontology (GO) enrichment analysis for TRAP-enriched genes. The GO categories are molecular function, biological process, and cellular component. The GO terms shown in the plots are the top ten most significantly enriched term for each list (non-redundant shown; Benjamini-Hochberg, p<0.05). The size of each dot indicates the number of genes (count) and the color indicates the p value (p.adjust). The GO information is in **Supplemental Table 4** and gene lists are in **Supplemental Table 5**.

To determine if there is overlap between genes that have *fru P1* TRAP-enriched expression in the gene lists from the different stages and sexes, we generated an UpSet plot (65), which is conceptually similar to a Venn diagram, but can accommodate a large number of conditions (**Figure 4B**). The histogram at the top of the UpSet plot shows the number of genes with overlap in the gene sets (**Figure 4B**). The dots beneath each bar show the gene sets that have the overlapping genes (**Figure 4B**). For example, the 10-12-day male and female *fru P1* TRAP-enriched data sets have a large number of genes that overlap (1,274 genes), as expected given the overlap shown in the Venn diagram (3,119 genes, **Figure 4A**). A similar result is shown for the 48hr APF TRAP-enriched data, with a large number of genes overlapping in the male and female *fru P1* TRAP-enriched genes (1,057 genes). The observation of sex-shared TRAP-enriched genes suggests that expression of stage-specific genes is important, in both males and females.

We conducted gene ontology (GO) term analysis on the *fru P1* TRAP-enriched genes to determine if there were genes with known functions overrepresented in the lists, examining the top ten GO terms for each list (**Figure 4E**, for all GO enrichment data see **Supplemental Table 4**). We examined GO term categories: ‘molecular function’, ‘biological process’ and ‘cellular component’. Given there is overlap across all the gene lists, many of the GO categories are shared and include several expected of neurons, though there are some unique categories for each time point (**Figure 4E**). For ‘molecular function’, the lists from the 1-day time points have several unique categories including ‘neuropeptide receptor binding’ and ‘hormone activity’ terms, and fewer shared categories. The 10-12 day male and female, and 48 hr male *fru P1*-enriched lists have chromatin binding as significant. For ‘biological process’ and ‘cellular component’, the lists from the adult samples, have ‘metabolic process’ categories and ‘mitochondrial’ categories that are unique. We also note that the 48hr APF and 10-12 day TRAP-enriched gene sets are enriched with genes that encode Immunoglobulin domain containing proteins/cell adhesion molecules (IgSFs), as we previously reported for 1-day adults (42). We have also validated that the Dpr/DIP IgSFs, a subset of these cell adhesion molecules, play a role *fru P1* neurons, further validating the data sets presented here (66).

We next examined the genes with sex-biased expression in TRAP samples, which were identified by comparing expression between the male and female TRAP samples. Given that the analysis was done on the exon level, we can find genes for which one exon may be male-biased and another female-biased, so the gene would be considered to have a transcript isoform with sex-biased expression in each sex (**Supplemental Table 6**). For example, at the 48hr APF stage there are 43 genes that have transcript isoforms with sex-biased expression in each sex, including *Sxl*, which is known to have male- and female-specific transcript isoforms (reviewed in 67). Additionally, we find a larger number of male-biased genes at 48hr APF (~1.4 fold), which is similar to what we reported for 1-day (~4.0 fold) (42), but at 10-12 there are fewer genes with sex-biased expression and more in females (~1.3 fold) (**Supplemental Figure 5C**). Further, more genes overlap between the 48hr APF and 1-day time points, in males and females, which might be expected given the small number of genes with sex-biased expression at 10-12 day (**Supplemental Figure 5D-F**). We conducted GO term analysis on the *fru P1* TRAP-biased genes to determine if there were genes with known functions overrepresented in the lists and show the top ten GO terms for each list (**Supplemental Figure 5E**, for all GO enrichment see **Supplemental Table 4**). Overall, the 10-12 day TRAP-biased gene lists have more unique GO terms, with several categories of genes expected to underlie adult behavior. Overall, this analysis demonstrates the importance of performing stage-specific analyses, given the differences in genes observed at each time point.

### Hierarchical clustering analysis of chromatin modifications for genes with mRNAs that are enriched in TRAP gene expression data sets

We next determined the pattern of chromatin modifications for genes with *fru P1* and *elav* TRAP-enriched expression at each time point. We examined heat maps to see the patterns of chromatin modifications, generated with the deepTools algorithm (**Figure 5**), as above. The patterns of chromatin modifications are similar for the male TRAP- and female TRAP-enriched genes, within time point (**Figure 5A-F**), as was true when all genes were examined (**Figure 3**). Furthermore, within each time point, there is no pattern of histone modifications that would clearly predict higher fold-enriched expression of a gene in *fru P1* neurons, with average fold differences in expression indicated on the left of each heat map (**Figure 5 A-F**). However, unexpectedly, at the 48hr time point, genes in a cluster with average 2-fold or greater *fru P1* TRAP-enriched expression have less enrichment of activating chromatin modifications H3K27ac and H3K4me3 (**Figure 5A-B**). We observe similar results with the *elav* chromatin data, where *elav* TRAP-enriched genes (41) have similar histone modification patterns (**Supplemental Figure 6**), but no large differences in the patterns that would predict gene expression levels.

**Figure 5.**
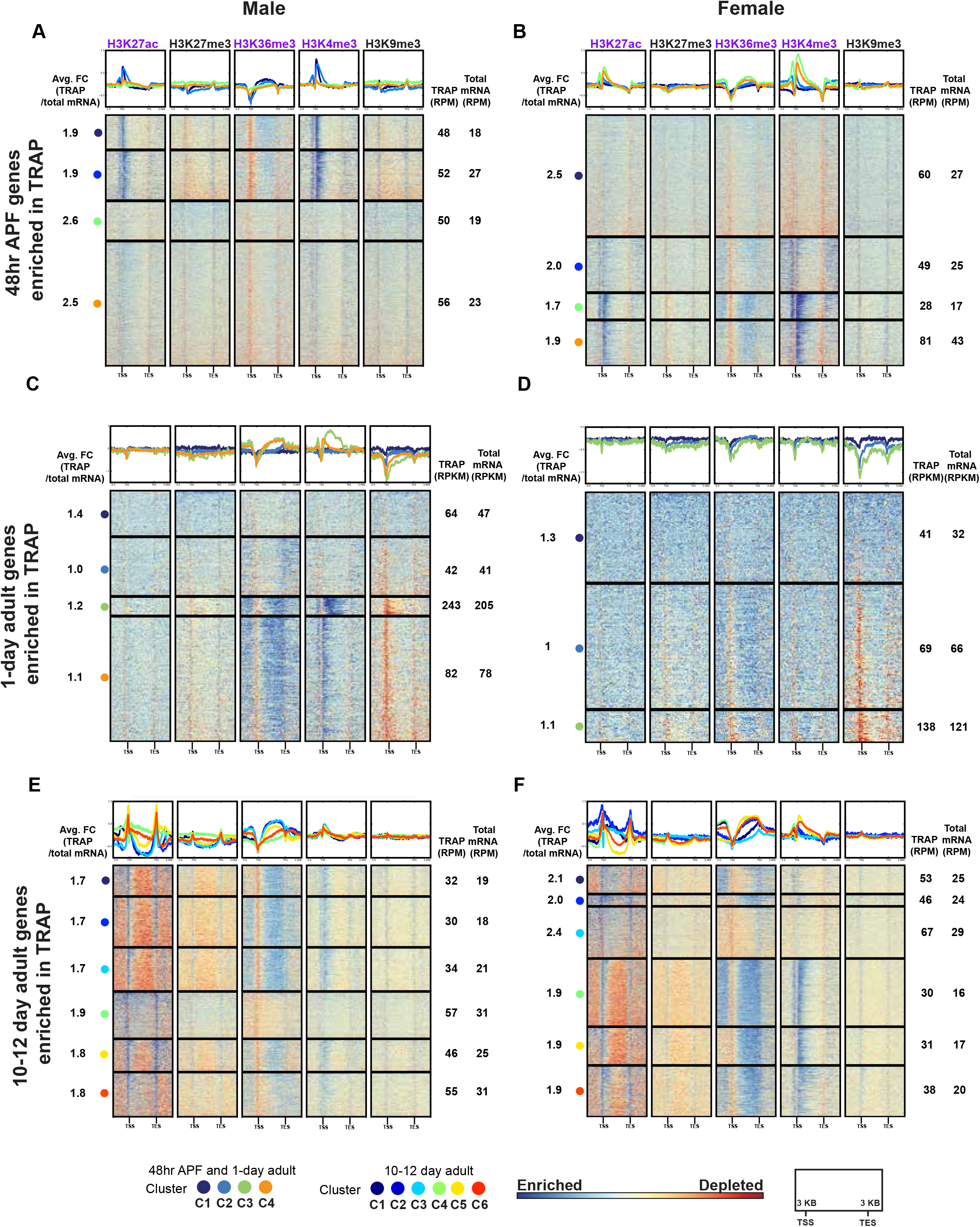
Hierarchical clustering of histone modification distributions for TRAP-enriched genes. Hierarchal clustering reveals shared combinations of histone modifications for TRAP-enriched genes, generated using deepTools. For detailed description of heatmaps see **Figure 3**. Heatmaps for *fru P1* chromatin data from 48hr APF **(A-B)**, 1-day **(C-D)**, and 10-12 day **(E-F)**. Gene lists for each cluster are provided in **Supplemental Table 2**. For each cluster of genes, the average fold-enrichment (Avg FC) of gene expression is indicated on the left (TRAP-enrichment), calculated at the exon level. For each cluster of genes, the average expression is indicated on the right for the TRAP and total mRNA data, calculated at the exon level. The 48hr APF and 10-12 day gene expression levels are provided as reads per million (RPM). The 1-day timepoint gene expression levels are provided as reads per kilobase per million (RPKM) (42). For TRAP expression data see **Supplemental Table 5**.

It appears that the chromatin modification patterns for TRAP-enriched genes indicate gene expression, but not levels of expression. To further address this, we examined the chromatin modification patterns for the genes with higher expression in total mRNA than TRAP-enriched mRNA. These genes have the same patterns of chromatin modifications as the TRAP-enriched genes (**Supplemental Figures 6** **and** **7**), for both *fru P1* and *elav* data sets. Overall, we find histone modification patterns associated with gene expression, but no patterns appear to be predictive of different levels of expression in *fru P1* or *elav* neurons. This result is also further confirmed by heatmaps made with genes that have detected expression in *fru P1* or *elav* neurons, rather than TRAP-enriched or total mRNA-enriched (**Supplemental Figure 8)**. This demonstrates that the histone modifications patterns may be predictive of gene expression in *fru P1* and *elav* neurons but are not predictive of gene expression levels.

One goal of this study was to examine how Fru^M^ impacts chromatin modifications, given a previous study showed that Fru^M^ recruits chromatin modifying enzymes (33). To assess if genes that are predicted to be directly bound by Fru^M^ have different chromatin modification enrichment profiles, we determined which clusters of *fru P1* TRAP-enriched genes have significant overlap with lists of genes that are predicted to be direct Fru^M^ targets (62, 63) (**Supplemental Table 3**). Based on visual inspection, the clusters with the more robust enrichment of activating chromatin modifications tend to be those that contain genes that are less likely to be direct Fru^M^ targets. Given that this is observed in both males and females heatmaps suggests the chromatin modification pattern differences in these clusters are not simply due to Fru^M^ binding activity.

### Identification of H3K27ac super-enhancers in chromatin from *fru P1* and *elav* neurons

Super-enhancers (SEs) are defined as regions of clustered enhancers and have been shown to regulate cell-type-specific gene expression (68). We identified SEs based on presence of clusters of H3K27ac peaks using the ROSE algorithm (68, 69). In brief, H3K27ac peaks located within a 12.5kb distance from each other were stitched together and plotted in an increasing order, based on their input-normalized H3K27ac ChIP-seq signal. Enhancers above the inflection point of the curve were defined as SEs (**Figure 6A-H**, **Supplemental Table 7**). Given the difficulty calling peaks on the male X-chromosome using MACS2, we perform the analysis on all chromosomes (**Figure 6**), and only autosomes (**Supplemental Figure 9**), and find overall the trends are similar, regardless of the X-chromosome SEs being included. For *fru P1* data sets, we find an increasing number of genes with SEs across developmental time in both males (61-267 SEs) and females (59-177 SEs), with the largest number identified in 10-12 day adult males (**Figure 6A-F**). We also identified genes with SEs in *elav* chromatin for males (552 SEs) and females (293 SEs) (**Figure 6G-H**).

**Figure 6.**
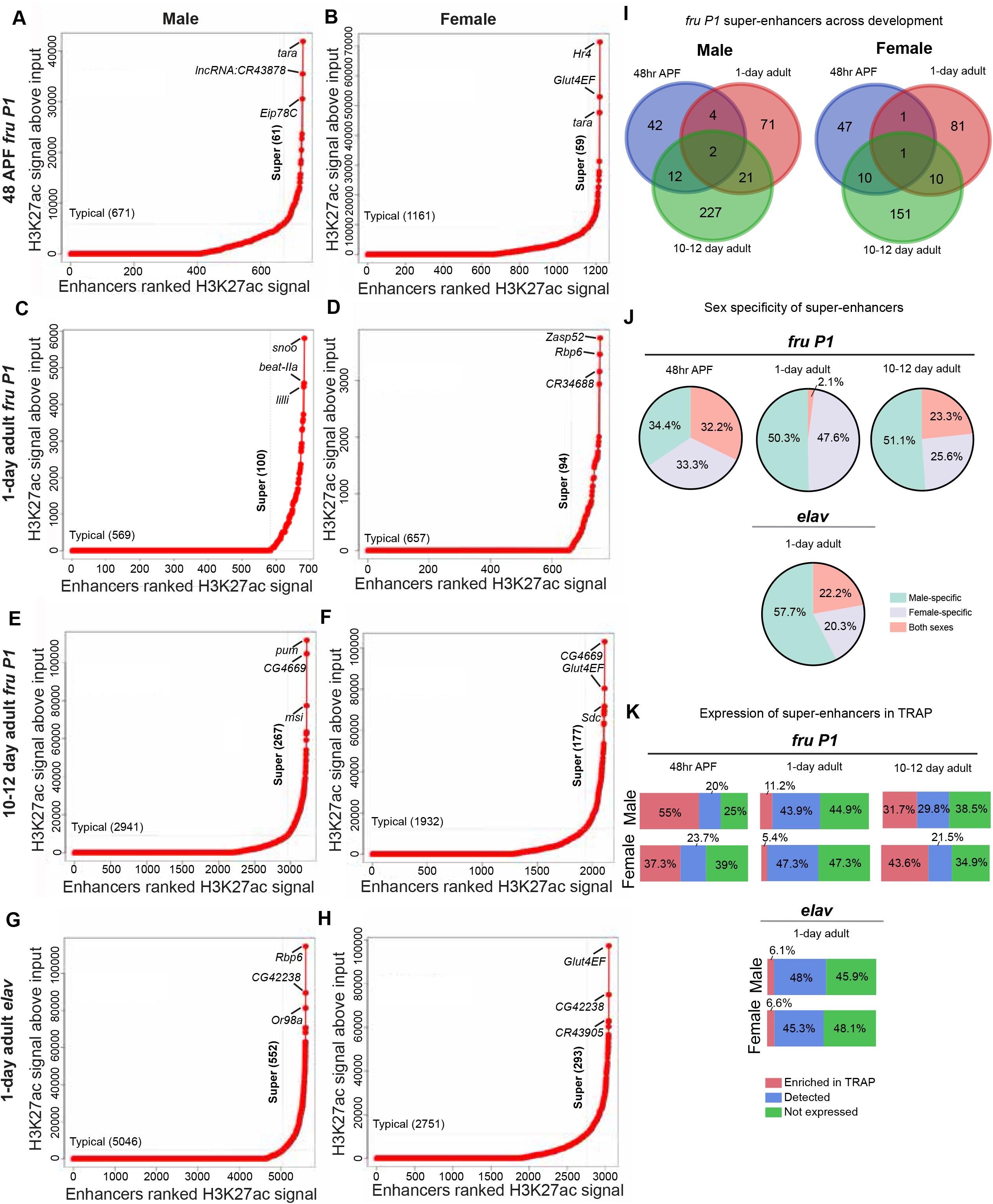
Super-enhancers identified based on H3K27ac peaks in *fru P1*- and *elav* neuron chromatin data sets. Super-enhancers (SEs) in *fru P1* **(A-F)** and *elav* (**G-H**) neuron chromatin data, identified using H3K27ac MACS2 peaks and the ROSE algorithm (68, 69). The ROSE algorithm was implemented with 12.5 kb regions stitched together and exclusion of regions +/− 2500bp to the TSS. The X-axis has the enhancers ranked by H3K27ac signal. The Y-axis indicates the H3K27ac signal above input. Dotted lines indicate the boundary between typical enhancers and SEs. The top 3 ranking SEs for each condition are indicated. **(I)** Venn diagrams showing the overlap between SE-containing genes across time points for *fru P1* chromatin data from males (left) and females (right). **(J)** Percentages of SE-containing genes for each stage and neuron type (*fru P1* or *elav*) that are male-specific, female-specific or occur in both sexes. **(K)** Percentages of SE-containing genes with detected or TRAP-enriched gene expression in *fru P1* or *elav* expression data sets.

When we compare SEs across time points, we find that the majority of the genes with SEs are stage-specific (**Figure 6I**), as shown in the Venn diagrams. Furthermore, genes with SEs also significantly overlap with lists of genes that are predicted to be direct Fru^M^ targets (62, 63) (**Supplemental Table 3**). The GO analysis of genes with SEs at 48hr APF reveals several enriched ‘neuron developmental’ terms, which are also enriched at 10-12 days. However, at 10-12 days we find additional enriched ‘adult behavior’ and ‘signaling’ terms (**Supplemental Table 4**). We next determined if there is sex-specificity of genes with SEs at each time point (**Figure 6J)**. Males and females have a similar of number of sex-specific genes with SEs at the early time points, but the 10-12 day time point has more genes with SEs that are unique to males (**Figure 6J)**. For the *elav* data sets, genes with SEs also have more male unique genes (**Figure 6J)**. However, we find that having a SE is not highly predictive of gene expression, when we compare to the *fru P1* and *elav* TRAP data sets from each time point (**Figure 6K**). Across the time points, we find a range of SE-containing genes that also have significantly enriched expression in the *fru P1* TRAP-enriched data sets (5.4%-55%; red squares in **Figure 6K**). This also true for genes with SEs in the *elav* data set, with only 6% of genes having enriched expression in the *elav* TRAP-enriched data set (41). Taken together, a gene containing a SE does not predict expression in the neurons at the time points examined.

### Candidate genes with super-enhancers and TRAP-enriched expression are expressed in *fru P1* neurons

Some genes were identified as having SEs, based on MACS2 H3K27ac peaks, and also having TRAP-enriched expression, so are a high confidence set of genes to determine if they overlap with *fru P1* expression. To determine co-expression, we used an intersectional genetic strategy, which is a modification of the Gal4/UAS system, with expression of a membrane-bound GFP marker indicating co-expression. The UAS-GFP reporter transgene has a stop cassette that can be removed by *fru P1* driven flippase expression (70). Once the stop cassette is removed the reporter gene expression is driven by a candidate-gene-specific-Gal4 driver (**Figure 7A**). We selected genes where there are stocks available that have *Gal4* transgenes (71, 72). We examined 10-12 day adult brains, where we can detect perdurance or concurrent expression of GFP. Nine out of 10 genes we screened exhibit expression in both sexes, consistent with our TRAP-enriched expression results (**Supplemental Table 6**). *glut4ef* only shows expression in female *fru P1* neurons, which might be due to the nearby female-specific SE identified at the 10-12 day stage that is not detected in males. However, *glut4ef* is TRAP-enriched in both males and females. This analysis also reveals sexual dimorphism in the co-expression patterns that could be due to differences in gene expression, but that remains to be determined. For example, seven lines with sex-specific SEs, have clear sexual dimorphisms apparent (*kibra*, *Eip75b*, *beat-IIa*, *rl*, *EcR*, *wbd*, and *glut4ef*). Though it should be noted that genes with SEs in both sexes also show sexual dimorphisms in their expression pattern (*gukh* and *tara*). Overall, all the candidate genes with SEs and TRAP-enriched expression had overlapping *fru P1* expression, demonstrating that these are a high confidence set of genes that are co-expressed with *fru P1*.

**Figure 7.**
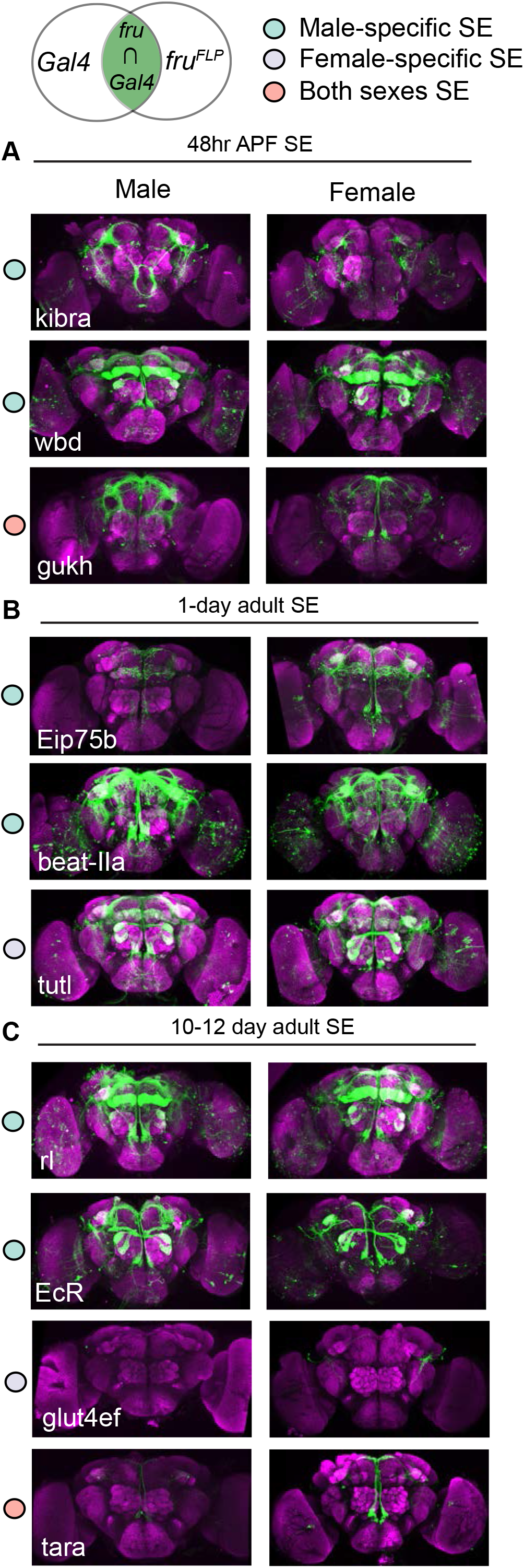
Co-expression of super-enhancers containing genes and *fru P1* in neurons. Super-enhancer containing genes were identified at each time point and their co-expression with *fru P1* was determined in 10-12 day adult brains. The top illustration shows the genetic intersectional approach where membrane bound GFP is only expression in neurons that co-express the candidate gene and *fru P1*. The top of each set of panels indicates the time point for which the super-enhancer was detected in the chromatin data sets: **(A)** 48hr APF stage, **(B)** 1-day adult, and **(C)** 10-12 day adult stage. The gene name is indicated in each panel, with intersecting expression patterns shown for male (left) and female (right) brains. Colored dots in legend indicate that the SE was identified in chromatin data from either male, female or both sexes.

## Discussion

Here, we describe Chromatag, a new molecular-genetic tool to perform cell-type-specific chromatin modification analyses that complements other existing approaches (reviewed in 34, 35). A benefit of the Chromatag approach is that FACs sorting is not needed, so the tissues are not perturbed before the chromatin fixation step and thus may better reflect the true chromatin state. Furthermore, this tool can be used with any Gal4 driver to examine any cell-type-specific chromatin modification or chromatin binding protein, for which there is a ChIP-grade antibody available, so is highly versatile. The wealth of Gal4 driver lines in *Drosophila* and the additional tools available to further restrict Gal4 expression opens up several possibilities to deepen our understanding of chromatin modifications and chromatin binding proteins. The results shown here demonstrate the approach is effective in small sets of neurons, at several developmental stages.

Our examination of five chromatin modifications in *fru P1* neurons, from three time points, revealed that there are large changes in chromatin modification profiles over developmental time and fewer sex-specific differences (**Figure 3**). These results show the importance of understanding the functional differences of chromatin modifications over developmental time, especially with respect to cell fate determination and adult physiological roles. One goal of this work was to determine if there is a chromatin modification pattern that might be predictive of gene expression in *fru P1* neurons. While the overall patterns of chromatin modifications are different for *fru P1* TRAP-enriched genes, as compared to all genes, there was not a highly predictive pattern for gene expression (**Figures 3** **and** **5**). This could be because the five chromatin modifications chosen are not the critical ones that would be indicative of gene expression in the cell-types we analyzed. Furthermore, given that we examined all *fru P1* neurons, we may miss chromatin modification patterns that are unique to a small set of cells, given the averaging of ChIP-seq signal across all neurons. Additionally, if a chromatin modification is transiently present, it may be difficult to detect. Finally, our TRAP-seq gene expression analyses are cell-type-specific but only includes the actively translated subset of mRNAs. Thus, the chromatin modification patterns may reflect overall gene expression that could be missed using TRAP RNA-seq. Given the advent of single-cell chromatin modification approaches, future studies can potentially resolve these issues.

The cell-type-specific TRAP gene expression studies add to the growing body of work that have identified genes important for *fru P1* function and contain a treasure trove to mine for further single-gene studies (42, 62, 63, 73–75). The new TRAP-seq data sets presented will aid in our understanding of how the potential for behaviors are established and maintained, by comparing the 48hr APF and 10-12 day adult data sets (**Figure 4** **and** **Supplemental Figure 5**). For example, we find many unique genes with TRAP-enriched expression at each time point, within sex. Furthermore, we find more male TRAP-enriched genes, compared to females, during 48hr APF, consistent with a similar observation at 1-day (42). At 10-12, this large skew of male TRAP-enriched genes is not observed. Additionally, the 48hr APF time point has the largest number of male-biased TRAP genes, suggesting a critical role for Fru^M^ during metamorphosis and early adult stages, which is also consistent with previous functional studies (28, 29, 64). Most GO terms identified at 48hr APF for TRAP-enriched genes are also found at 10-12 day, though there are several unique terms at 10-12 day. When we closely examined the GO enrichment for these genes we found that many of the of the terms are related to ‘chromatin’, ‘G protein-coupled receptor activities’ and ‘RNA regulation’ (**Figure 4C**, **Supplemental Table 4**). These results are consistent with findings of Fru^M^ and *fru P1* roles for adult-specific functions (30, 32). Altogether, this suggests that there are dynamic gene expression changes occurring in these neurons in both sexes, as well as sex-specific gene expression which may contribute to sex differences in neuronal physiology in adults to generate male and female reproductive behaviors.

While the goal of this study was to gain insight into *fru P1* regulation of genes, the results presented can also serve as a launching point to further understand the functional roles of chromatin modifications and SEs. For example, we find that SEs do not always predict gene expression, suggesting that in some cases the SEs may mark a gene as poised for expression. The SE activity that directs gene expression may depend on the presence of other activating modifications that are found in enhancer regions, such as H3K4me1 (38). Others have reported similar observations where computationally predicted enhancers are not necessarily linked to gene expression activity and show a dependence on transcription factor binding (76, 77). Furthermore, the Bachtrog laboratory also found that chromatin state did not account for sex-biased gene expression in studies performed at the larval stage (78). Given that Fru^M^ has been implicated in recruiting chromatin modifiers, further direct Fru^M^ occupancy studies at these stages may elucidate its role in regulating gene expression through histone modifications. The application of machine learning approaches may also reveal new insights about the role of chromatin modifications in directing gene expression. Additional studies using techniques such as targeting chromatin modifying enzymes to particular loci, using Cas9-based approaches, can address the functional roles of the modifications presented here (reviewed in 79, 80). Moreover, *Drosophila* reproductive behaviors are an excellent model for studies examining how chromatin modifications and gene expression underlie behavioral plasticity, using models like long-term memory of courtship rejection and female post-mating behaviors (9, reviewed in 81). The tools and the courtship model described here are a premiere system to go from high resolution genome-wide chromatin modifications and gene expression studies, to understand the functional impacts of these molecular phenotypes on neuroanatomy and ultimately behavior.

## Materials and Methods

### Chromotag fly constructs and strain generation

The H2B open reading frame was PCR amplified using high-fidelity Taq. The NsiI and SalI restriction enzyme sites were added to the 5’ and 3’ end, respectively, during PCR amplification, using the following oligonucleotides: atgcatgatATGCCTCCGAAAACTAGTGGAAAGGCAGCC and gtcgacTTATTTAGAGCTGGTGTACTTGGTGACAGCCTTGG. The resulting PCR product was Topo cloned (Invitrogen) and sequenced verified to ensure no PCR mutations were introduced. The H2B open reading frame was subsequently cloned into the p-UAS-3XFLAG-BLRP-mcherry-3’UTR plasmid for expression in *Drosophila*, using the Gal4/UAS system (plasmids provided by Paul Talbert and Steve Henikoff). The UAS-3XFLAG-BLRP-mcherry-H2B-3’UTR cassette was then cloned into the p2-Casp-UAS-BirA plasmid that contains the *E. coli* BirA gene under UAS control, P element ends and the mini-*white* gene (also provided by Paul Talbert and Steve Henikoff). This construct was injected into *Drosophila white* strains for P element transformation by Genetivision Inc.

### *Drosophila* strains used in genomic assays

Flies with Chromatag expressed in *fru P1* neurons or broadly expressed with a pan-neuronal *elav* driver were the respective genotypes: *w;P[w^+mC^*, *UAS-Gal4]/P[UAS-Chromatag]*; *fru P1-Gal4*/+ or *w;P[w^+mC^*, *UAS-Gal4]/P[UAS-Chromatag]*; *elav-Gal4*/+. Flies expressing the GFP-tagged variant of RpL10A in *fru P1* neurons were the genotype: *w/(w or* Y); *P[w*^+*mC*^, *UAS-Gal4]/P[w*^+*mC*^, *UAS-GFP*::*RpL10A]*; *fru P1-Gal4*/+(41). The *elav-Gal4* strain was obtained from the Bloomington stock center. Flies were reared on standard cornmeal food at 25°C on a 12-h light/12-h dark cycle. List of fly stocks used in this study are reported in **Supplemental Table 8**.

### *Drosophila* tissue collection

For cell-type-specific ChIP-seq and TRAP-seq, approximately 1,000 sex-sorted flies were used. The time points were 48hr after pupal formation (APF, +/− 1hr), 16-24hrs post-eclosion (TRAP-seq see 42), or 10-12 days post-eclosion. For each ChIP-seq condition, 3-4 independent biological replicates were used. For each TRAP-seq condition, 4-5 biological replicates were used. For the 48hr APF time point, flies were collected and sex sorted at the white prepupal stage and aged for two-days. For the 1-day time point, 0-8hrs post-eclosion flies were collected. For the 10-12 day time point, bottles were cleared and adults were collected within two-days. For the adult time points, flies were aged to either 16-24hrs after eclosion or 10-12 days after eclosion and snapped frozen without additional CO_2_ anesthesia. All flies were frozen at 1-2 hours after incubator lights turn on. All frozen animals were stored at −80°C until they were processed for downstream experiments.

### Translating ribosome affinity purification

Translated mRNA purification was performed as described in Newell *et al.* 2016 with the following modifications: at the last step, prior to library preparation the 40μl of lysate supernatant (input, total mRNA) and TRAP (IP) fractions were resuspended with RA1 lysis buffer to a total volume of 200μl for one additional column-based purification (Machery-Nagel Nucleospin RNA clean-up, product 7404948), using DNAse.

### Sequential ChIP

Frozen adult animals were used and heads were isolated. The heads were separated from their bodies by vortexing frozen animals. The heads were sifted through liquid nitrogen-cooled mesh sieves #25 and #40 (with heads retained in the #40 sieve). Frozen adult heads or frozen whole body pupae were dounce homogenized in buffer A1 (60mM KCL, 15mM NaCl, 4mM MgCl_2_, 15mM HEPES (pH 7.6), 0.5% TritonX-100, 0.5mM DTT, 10mM sodium butyrate, and Roche cOmplete Protease Inhibitor cocktail EDTA-free, product 5056489001), containing methanol-free 1% formaldehyde solution (Pierce, product 28906) for crosslinking at room temperature for 10 min. The crosslinking reaction was quenched using 2.5M Glycine and incubated for 5 minutes at room temperature.

Cross-linked homogenates were centrifuged at 4000g for 5 minutes at 4°C, and the nuclear pellets were resuspended in buffer A1, and centrifuged at 4000g for 5 minutes at 4°C. The nuclear pellet was then washed by resuspension in buffer A1 and centrifuged as described above two more times. The nuclear pellet was then resuspended in 1x Lysis Buffer without SDS (140mM NaCl, 15mM HEPES (pH 7.6), 1mM EDTA (pH 8.0), 0.5mM EGTA, 1% TritonX-100, 0.5mM DTT, 0.1% sodium deoxycholate, 10mM sodium butyrate, and Roche cOmplete Protease Inhibitor Cocktail EDTA-free) to remove all traces of A1 buffer before the next lysis step. The washed nuclear pellet was resuspended in Lysis buffer containing 1% SDS and 0.5% N-Laurosylsarcosine and incubated at 4°C, rotating for 1 hour. Nuclear lysates were quickly spun down for 10 seconds at 2000g and supernatant transferred to sonication tubes (Covaris; product 520130) for shearing. Chromatin was sheared for 25 minutes using a Covaris E220 (duty cycle: 5.0, peak incident power: 140, cycles per burst: 200). The sheared chromatin was transferred to microcentrifuge tubes and rotated for 10 min. at 4°C and then centrifuged at 18,000g for 5 min. at 16 °C. Sheared chromatin was incubated with 40μl of magnetic protein A/G beads (Pierce, product 88803) and 5μg of each respective antibody at 4°C overnight. The magnetic protein A/G beads were then washed four times with lysis buffer wash (140mM NaCl, 15mM HEPES (pH 7.6), 1mM EDTA (pH 8.0), 0.5mM EGTA, 1% TritonX-100, 0.05% SDS, 0.5mM DTT, 0.1% sodium deoxycholate, 10mM sodium butyrate, and Roche cOmplete Protease Inhibitor Cocktail EDTA-free) and twice with TE (10mM Tris-HCL (pH 8.0), 1mM EDTA) for 10 minutes each, rotating at 4 °C. Beads were incubated in ChIP elution buffer (100mM NaHCO_3_, 80mM NaOH, 1% SDS) at 37°C, rotating for 30 minutes to elute chromatin. The eluate was diluted 1:10 with Lysis buffer without SDS and incubated on magnetic streptavidin beads (Invitrogen Dynabeads MyOne Streptavidin C1) for 4hrs-overnight rotating at 4°C. The beads were washed as above and chromatin was reverse crosslinked on beads in ChIP elution buffer (with 0.04mg/ml proteinase K, 0.01mg/ml RNaseA) at 65 °C overnight. DNA was purified with a DNA purification and concentration kit (Machery-Nagel, product 740609).

### Illumina sequencing and library preparation

The isolated total RNA for TRAP libraries underwent poly(A)^+^ selection using the NEBNext Poly(A) mRNA isolation module (NEB, product E7490S). 48hr APF TRAP libraries were prepared with the NEBNext Ultra II Directional RNA Library Prep Kit for Illumina (NEB, product E7760S) and 10-12 day adult TRAP libraries were prepared with the NEBNext Ultra RNA library Prep Kit for Illumina (NEB, product E7530S). ChIP-seq libraries were prepared using the NEBNext ChIP-seq Library Prep Master Mix Set for Illumina (NEB, product E6240). Libraries were size selected using AMPure XP beads (Beckman) and clean up steps were performed using the Machery-Nagel PCR cleanup kit (Machery-Nagel, product 740609). For ChIP-seq and 48hr APF TRAP-seq, a final clean-up using AMpure XP beads was performed according to kit protocol to remove excess primers. Samples were pooled and sequenced on the Illumina HiSeq 2500 platform with 100-bp single end reads for 10-12 day adult TRAP libraries and 50-bp single end reads for ChIP-seq libraries. 48hr APF TRAP libraries were sequenced on the Illumina NovaSeq 6000 with 100-bp single end reads on an S1 flowcell.

### Sequencing read mapping

Read quality was assessed for each ChIP-seq and RNA-seq sequencing library with FastQC (v. 0.11.3). ChIP-seq reads were pre-processed using Trimmomatic’s ILLUMINACLIP (v. 0.33) option to remove Illumina adaptor sequences (83). Pre-processed ChIP-seq reads were aligned to the *D. melanogaster* dm6 UCSC genome assembly (84), using Bowtie2 (v. 2.2.5), allowing up to 6 mismatches (85). Only uniquely aligned reads were considered and individual replicate and pooled replicate (all replicates) BAM files were generated with SAMtools (v. 1.2) for downstream analyses (86). Sequencing coverage for pooled replicates for histone H3 and all histone modifications in this study were calculated using the pileup.sh function in BBMap on the uniquely mapped reads. RNA-seq reads were aligned using STAR (v. 2.7.2a) to the same genome assembly as previous (87).

### ChIP-seq peak calling and analysis

Regions of significant enrichment (peaks) for each histone modification (H3K27ac, H3K36me3, H3K4me3, H3K9me3, H3K27me3), at each time point (48hr APF, 1-day adult, 10-12 day adult), for each sex in *fru P1* or *elav* neurons, relative to sex and stage-specific input chromatin library controls for individual and the pooled replicates, were identified using the Model-based Analysis of ChIP-Seq, MACS2 (v.2.1.0) (55). For each replicate, the peaks can be found in the GEO upload. For the pooled and consensus peaks see **Supplemental Table 1**. We performed “narrow” peak calling for histone modifications with more compact enrichment patterns (H3K27ac and H3K4me3) and the “broad” peak option for modifications with diffuse enrichment (H3K36me3, H3K9me3, and H3K27me3) using the following: “callpeak” parameters: --gsize dm -p 1e-2 --call-summits --fix-bimodal (88).

In an effort to resolve more peaks on the male X-chromosome for all data sets we tried several modifications to our “callpeak” parameters by systematically adjusting --extsize, --bw, -q, and -p, --mfold, and --lambda (data not shown). We also performed SICER peak calling, implemented through epic2, as an alternative peak calling approach. However, none of these changes to MACS2 or use of SICER achieved a sizeable increase in peaks called on the male X-chromosome. Consensus peak sets among biological replicates were called using DiffBind requiring a peak to be present in 50% or more replicates (89). MACS2 peak lists were annotated using the “annotatePeak” function in CHIPseeker, using default settings using the TxDb.Dmelanogaster.UCSC.dm6.ensGene database (v. 3.8) (peaks listed in **Supplemental Table 1**) (90). Annotations include a genomic feature assignment which indicates whether a peak is located in a TSS, exon, 5’ UTR, 3’ UTR, intronic, or intergenic region which was visualized for each experimental condition using the “plotAnnoBar function”.

### ChIP-seq correlation, visualization, and clustering analysis

Input normalized BigWig files for all histone modifications, replicates, and conditions were obtained using deepTools “bamCompare” function to normalize to input ChIP-seq read counts across the entire genome, over default 50bp bins with a log2 ratio summary of signal for each bin reported (52). Using these normalized BigWig files, each experimental condition was compared using the “multiBigWigSummary” module to score ChIP-seq signal over 10kb windows (bins) across the genome and compressed into a matrix. These matrices were utilized in the “plotCorrelation” module for correlation analysis (Spearman correlation) to calculate similarity across histone modifications for both sexes, time points, and neuron types in our study (**Supplemental Figure 3**). To visualize normalized ChIP-seq signal over specified genomic regions of +/− 3kb of the TSS and TES, we used deepTools “computeMatrix” function (--beforeRegionStartLength 3000 -regionbodylength 5000 --afterRegionStartlength 3000) and the “plotHeatmap” function to perform hierarchical clustering (ward linkage) and heatmaps were generated. Visual inspection of multiple clustering outputs was performed to select the optimal number of clusters in each analysis.

### RNA-seq analysis

Exon-level count tables were generated in FeatureCounts, for all replicates, using uniquely mapped reads for male and female *fru P1* TRAP and total mRNA for both 48hr APF and 10-12 day adult data sets (91). Exons were only retained in the analysis if they met our filtering criteria: exons that had counts above 2 counts-per-million (CPM) in 60% of the replicates for each pair-wise analysis were retained. The pairwise comparisions are: 1) male TRAP vs. male total mRNA ; 2) female TRAP vs. female total mRNA ; 3) male total mRNA vs. female total mRNA; and 4) male TRAP vs. female TRAP. Filtered read counts were TMM-normalized and differential expression testing was performed on the exon level using edgeR’s Fisher’s exact testing for each pairwise comparison (92). Significant differentially expressed exons were those with an FDR of <0.2 (Benjamini and Hochberg FDR-adjusted p value). All exons and associated FDR-adjusted p values are listed in **Supplemental Tables 5 and 6**).

### Gene Ontology (GO) and pathway enrichment analyses

GO analysis and pathway enrichment analysis were conducted in R, using the clusterProfiler package, using a hypergeometric test for enrichment (93). Additional GO analyses, protein domain, and pathway enrichment were carried out in Flymine (60).

### Immunohistochemistry and microscopy

Adult and 48hr APF brains were dissected in 1x Phosphate Buffered Saline (PBS; 140 mM NaCl, 10 mM phosphate buffer, and 3 mM KCl, pH 7.4) and fixed in 4% paraformaldehyde (Electron Microscopy Sciences, product 15713) in PBS at room temperature, for 20 minutes for adult brains, or 45 minutes for 48hr pupal brains and ventral nerve cords. Next, the brains were rinsed briefly 4 times with PBS and then washed three times with TNT (0.1M Tris-HCL, 0.3M NaCl, 0.5% Triton X-100), for 15 minutes for the first wash and 5 minutes for the two subsequent washes. The brains were blocked in Image-iT FX Signal Enhancer (Thermo Fisher) for 25 minutes and washed twice with TNT for 5 minutes each. Brains were incubated in TNT with primary antibody overnight at 4°C. After primary antibody was removed, the brains were washed six times with TNT for 5 minutes per wash. Next, the brains were incubated in TNT with secondary antibody for two hours at room temperature. The brains were then washed six times for 5 minutes in TNT, before being mounted in Vectashield mounting medium (Vector Laboratories, H-1000) on glass slides in Secureseal^™^ Image Spacers (Electron Microscopy Services) and covered with a no. 1.5 glass coverslip. Confocal microscopy was performed on a Zeiss LSM 700 system, with 20x objective and bidirectional scanning. The interval of each slice was set as 1.0 μm. Zeiss Zen software (Black edition, 2012) was used to make adjustments to laser power and detector gain to enhance the signal to noise ratio.

### Antibodies

The antibodies used for ChIP were: H3 (Abcam ab1791), H3K27ac (Abcam ab4729), H3K27me3 (Millipore 07-499), H3K36me3 (Abcam ab9050), H3K4me3 (Abcam ab8580), and H3K9me3 (Abcam ab8898), and were validated by the company for specificity. The following primary antibodies were used for immunofluorescence: mouse anti-Nc82 (1∶20; Developmental Studies Hybridoma Bank) and rat anti-Fru^M^ (1:200)(18). The secondary antibodies used for immunofluorescence were Alexa Fluor goat anti-rat 488 (1:1000), goat anti-mouse 633 (1:500), rabbit anti-GFP 488 conjugate (1:600), and streptavidin 488 conjugate (1:4000) (Thermo Fisher).

### Behavioral assays

For courtship assays, all males were collected 0-6 hours post-eclosion and aged individually for 4-7 days at 25°C on a 12-h light/12-h dark cycle. The single male flies were either paired with a 4-7 day old *w; Canton S* (*CS*) virgin female (male-female) or male (male-male). Females were aged in groups of ~10 per vial. The assays were performed in a 10mm courtship chamber. Courtship activity was recorded for 10 min or until successful copulation at five-nine hours after fly incubator lights on, at 25°C (n=12). Recordings were analyzed using Noldus software. Courtship Index (CI) and Wing Extension Index (WEI) were calculated by dividing the time the experimental male fly spent courting or unilaterally extending his wing toward the target female by the total observation time. JMP Pro 14 was used to conduct statistical analyses and statistical significance (p<0.05) was calculated using a nonparametric Wilcoxon test (Mann-Whitney U test) for comparing CI and WEI of experimental males. The experimental male genotype is: *w; P[w*^+*mC*^, *UAS-Gal4]/P[UAS-Chromotag]*; *fru P1-Gal4*/+. The single transgene controls are: *w; P[w*^+*mC*^, *UAS-Gal4]*/+; *fru P1-Gal4*/+, and *w; P[UAS-Chromotag]*.

For locomotor activity, male and female flies of genotypes: *CS*, *w; P[w*^+*mC*^, *UAS-Gal4]*/+; *fru P1-Gal4*/+, and *w; P[UAS-Chromotag]*, and *w; P[w*^+*mC*^, *UAS-Gal4]/P[UAS-Chromotag]*; *fru P1-Gal4*/+ were collected 0-6 hours post-eclosion, briefly anesthetized with CO_2_, loaded into glass tubes containing standard laboratory food, and tubes were mounted in *Drosophila* Activity Monitor (DAM) boards (Trikinetics, Waltham, MA). Data was collected in 5 minute bins and experiments were carried out at 25°C with a 12-h light/12-h dark cycle for 8 days. The first 24hrs of data was removed from the analysis, to recover CO_2_ and acclimate to the DAM system. Data analysis was performed with ShinyR-DAM (94) and R script analysis in R studio.

### Identification of enhancers and super-enhancers

Super-enhancer (SE) analyses were conducted using the ROSE algorithm: (https://bitbucket.org/young_computation/rose) (68, 69). Briefly, MACS2 H3K27ac peaks were stitched together, if within 12.5 kb of one another, for pooled replicates for each experimental condition. Regions +/− 2500bp to the TSS were excluded to remove promotor bias. The ROSE algorithm then determined the H3K27ac intensity inflexion point to determine typical enhancers versus super-enhancers (for all enhancers see **Supplemental Table 7**), as compared to input ChIP signal (68, 69). Genomic regions containing enhancers or super-enhancers were annotated using the default “annotatePeak” function in CHIPseeker (90). H3K27ac SE-containing genes were compared to *fru P1* TRAP-enriched genes (defined above) and detected genes. A gene is considered detected if an exon passed our initial filter for the TRAP data sets but did not show significantly higher expression than total mRNA (**Supplemental Table 5**). This criteria was also used for *elav* TRAP, but this analysis was conducted on the gene level instead of the exon level (41).

### Gene overlap and enrichment analysis

Venn diagrams to visualize overlapping genes were generated using online tools at http://bioinformatics.psb.ugent.be/webtools/Venn/ and size proportional Venn diagrams were made using BioVenn (95). UpSet plots were generated using the UpSetR package (65). Enrichment between gene lists was analyzed using the GeneOverlap R Package, which performs Fisher’s exact tests to determine whether enrichment is significant given genome size and reports a Jaccard index overlap and odds ratio (61). All contingency tables reported in **Supplemental Table 3**

## Data availability

The Gene Expression Omnibus accession number for all ChIP and RNA-seq the data is: GSE153901

## Acknowledgements

The work was supported by NIH grants awarded to MNA: R01GM073039, R01GM116998. This work was also supported by funds from the Biomedical Sciences Department, College of Medicine, Florida State University. We are grateful for the support. We thank Steve Henikoff and Paul Talbert for plasmid reagents and helpful advice. We appreciate that colleagues provided Drosophila stocks. Stocks were also obtained from the Bloomington Drosophila Stock Center (NIH P40OD018537). Antibodies used in this study were obtained from the Developmental Studies Hybridoma Bank, created by the NICHD of the NIH and maintained at The University of Iowa, Department of Biology, Iowa City, IA 52242. We appreciate comments on the manuscript from Catherina Artikis. We thank Brandon Bowers for technical assistance. We thank the Biomedical Sciences Translation Core Staff for technical support.

---

**Supplemental Figure 1.**
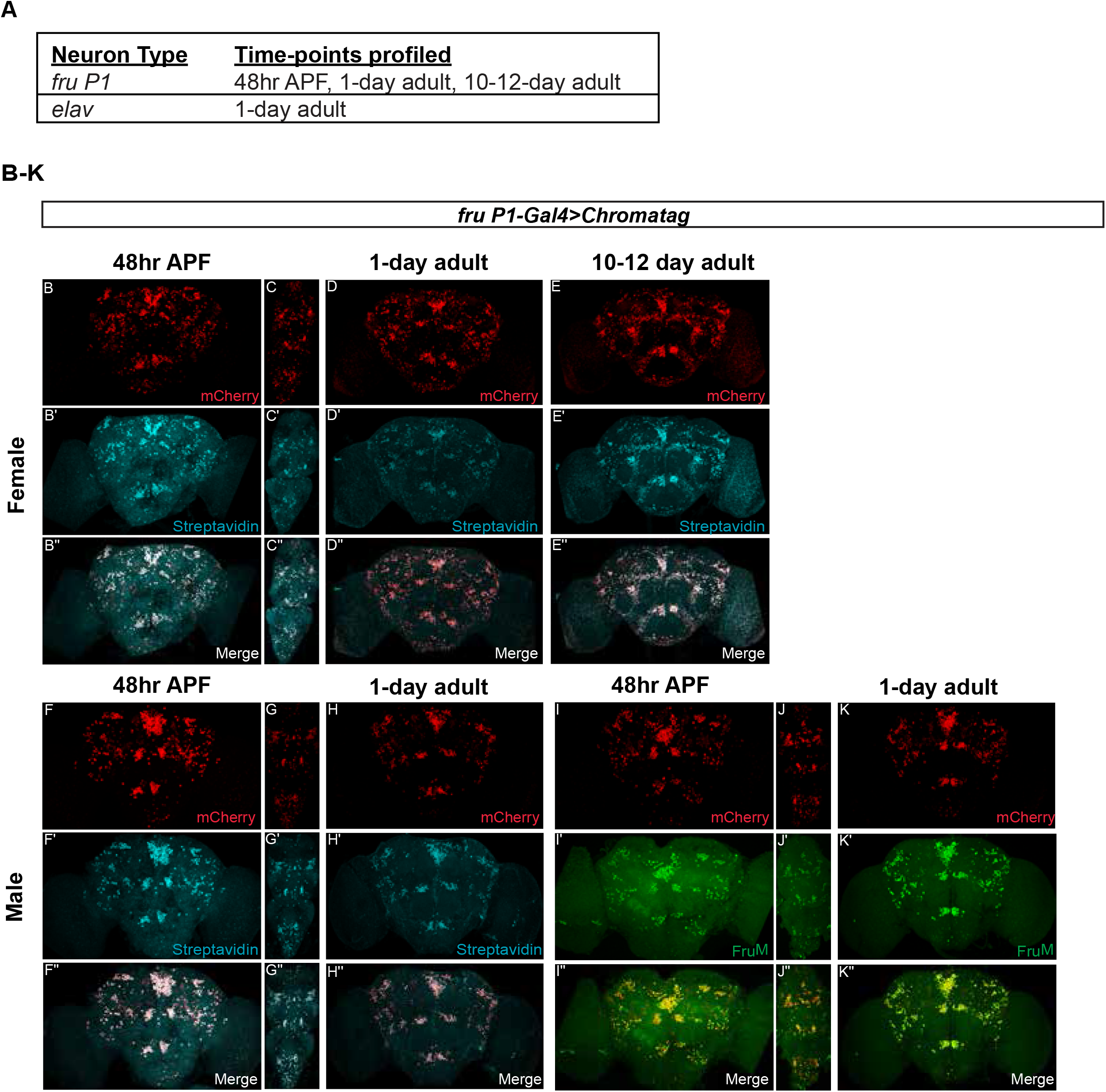
Experimental overview and visualization of *UAS-Chromatag* expression in *fru P1* neurons across time points. (**A**) Sequential ChIP-seq was performed in *fru P1* neurons from both sexes, at three time points: 48 hours after pupal formation (APF), 1-day adults (heads), and 10-12 day adults (heads) as well as in *elav-Gal4* expressing neurons in 1-day adult heads of both sexes. (**B-K**) Confocal maximum projections show *fru P1-Gal4* driven expression of *Chromatag*, which results in production of a tagged H2B variant that can be biotinylated. The H2B variant is produced and detected at all time points and in both sexes, used in this study (also see **Figure 1**). Brains and ventral nerve cords are shown in alternate columns. Biotinylated H2B is detected by fluor-conjugated streptavidin (cyan). Fru^M^ is detected by immunostaining (green), using an anti-Fru^M^ antibody. mCherry tagged-H2B is visualized by genetically encoded mCherry.

**Supplemental Figure 2.**
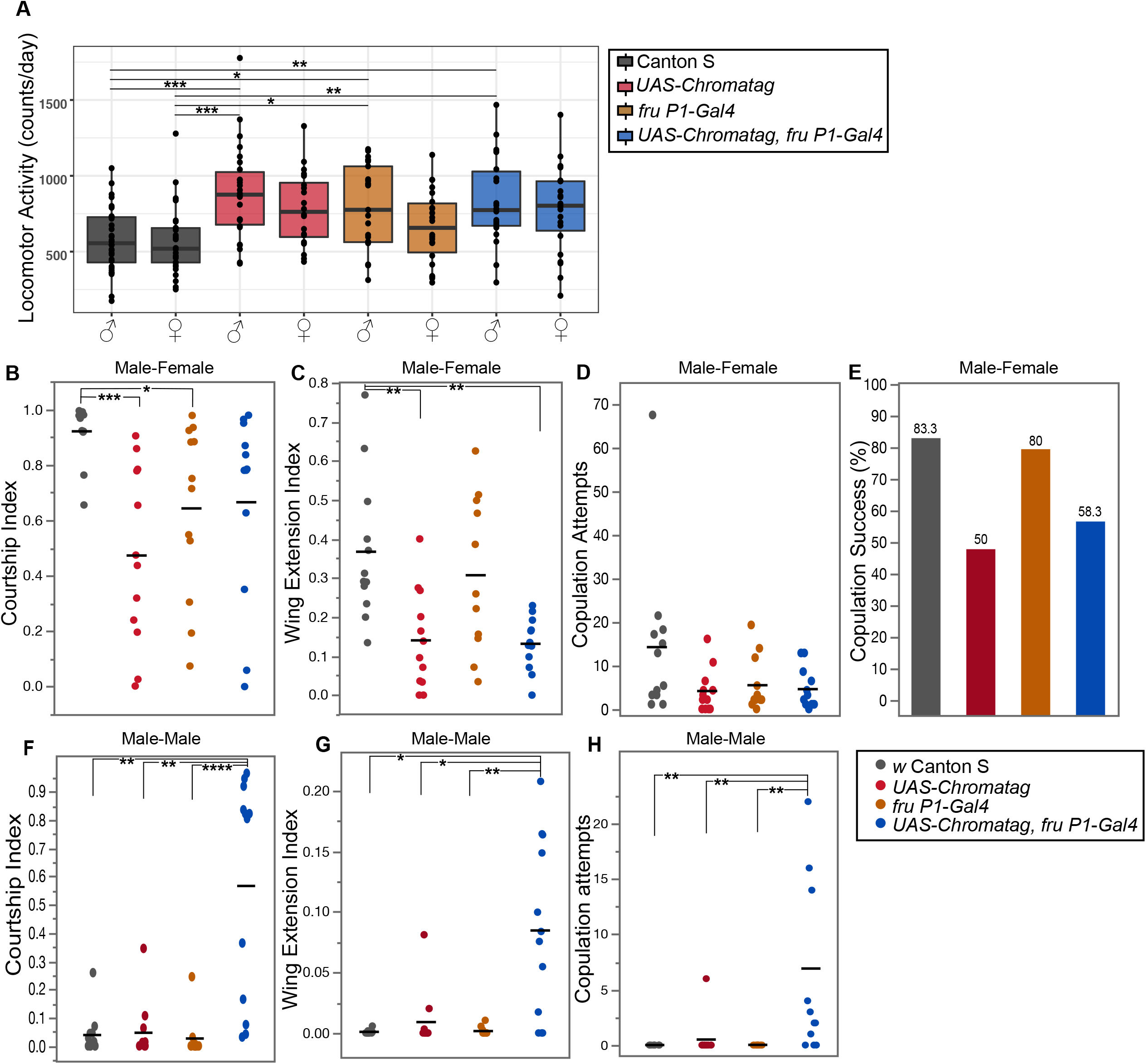
Locomotor activity and courtship behavior of males expressing *Chromatag* in *fru P1* neurons. **(A)** Locomotor activity of males and females (n=20-32 individuals per genotype). The average counts of beam crossing per individual, per day, is shown in boxplots, where upper and lower hinges correspond to the 25th and 75th percentiles (data collected using Drosophila Activity Monitors; Trikinetics). Activity data was analyzed by one-way ANOVA. Tukey HSD post-hoc tests were performed for each pairwise comparisons and significant differences are shown. **(B-H)** Male courtship toward a female (**B-E**), or male (**F-H**). The behavioral indices are the total time the male spends performing the courtship behavior/total video time, or the time until copulation occurs. (**B**) Male courtship index. (**C**) Wing extension index. (**D**) The number of copulation attempts toward the target female. (**E**) Percentage of males that successfully copulate with target females. (**F-H**) Male courtship toward *white Canton S* target male. (**F**) Male courtship index. (**G**) Wing extension index. (**H**) The number of copulation attempts toward the target male. (**B-H**) Horizontal bars show the mean (n=12 males). Statistics performed for all pair-wise comparisons. Kruskal-Wallis ANOVA test with Dunn’s post-hoc correction used in tests with courtship and wing extension index data. One-way ANOVA with Tukey post-hoc analysis performed on copulation attempts data. Significance levels are indicated: p<0.05 *, p<0.01 **, p<0.001 ***, p<0.0001 ****.

**Supplemental Figure 3.**
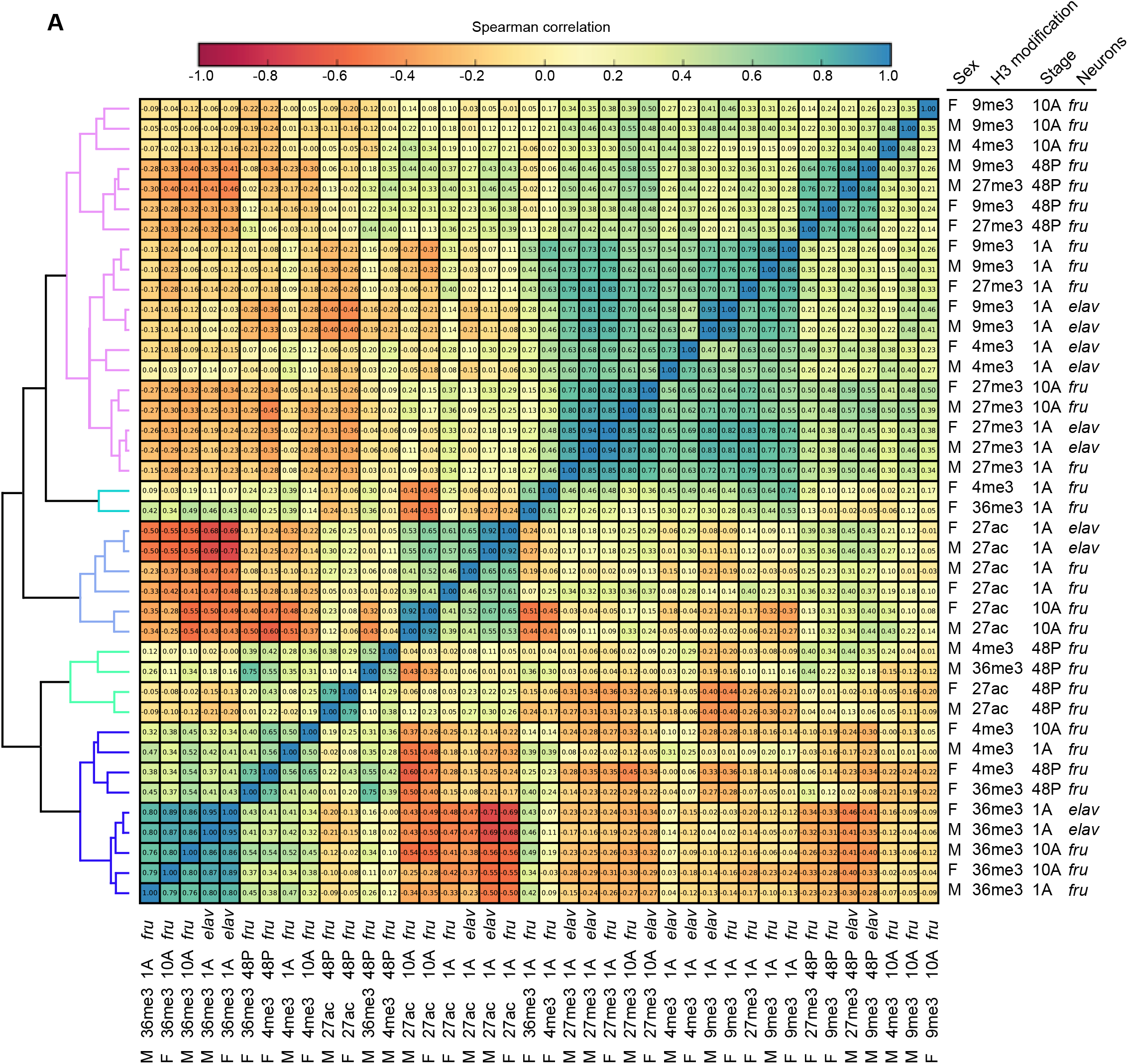
Spearman correlation of Chromatag-ChIP data sets. (**A**) The positional correlation of genome-wide ChIP-seq signals from all the data sets, comparing 10kb bins, using unsupervised hierarchical clustering is shown. The level of positive or negative correlation is indicated by color (see scale bar). The dendrogram on the left indicates which samples read positions are most similar to each other. Additional color coding by groups with positive correlation is presented. For each chromatin data set, the sex, H3 modification, time point and neuronal cell type is indicated. This analysis was performed using the pooled replicates for each data set (n=3-4).

**Supplemental Figure 4.**
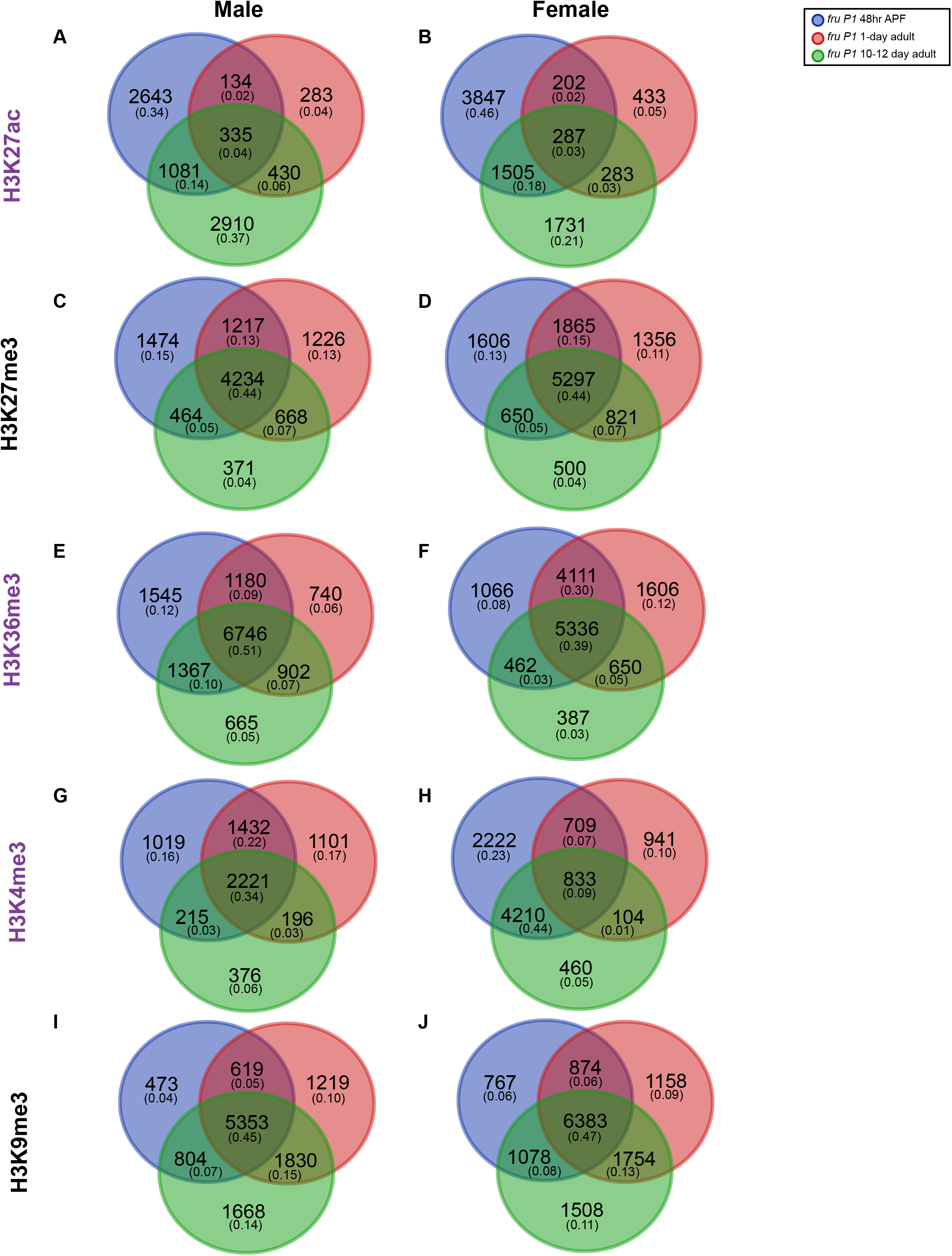
Overlap of genes containing MACS2 peaks across time points in *fru P1* neurons. **(A-J)** Venn diagrams comparing genes that have at least one MACS2 peak for each chromatin modification, across time points and within sex. For each Venn diagram category the number of genes and the proportion of the total genes (in parentheses) in each panel is shown. The histone modification is indicated on the left (activating in purple, repressive in black). The legend for each time point is on the top left and male and female data sets are indicated at the top. All MACS2 peaks were called on pooled biological replicates (n=3-4) and identified as enriched relative to matched input controls. MACS2 peaks are listed in **Supplemental Table 1**.

**Supplemental Figure 5.**
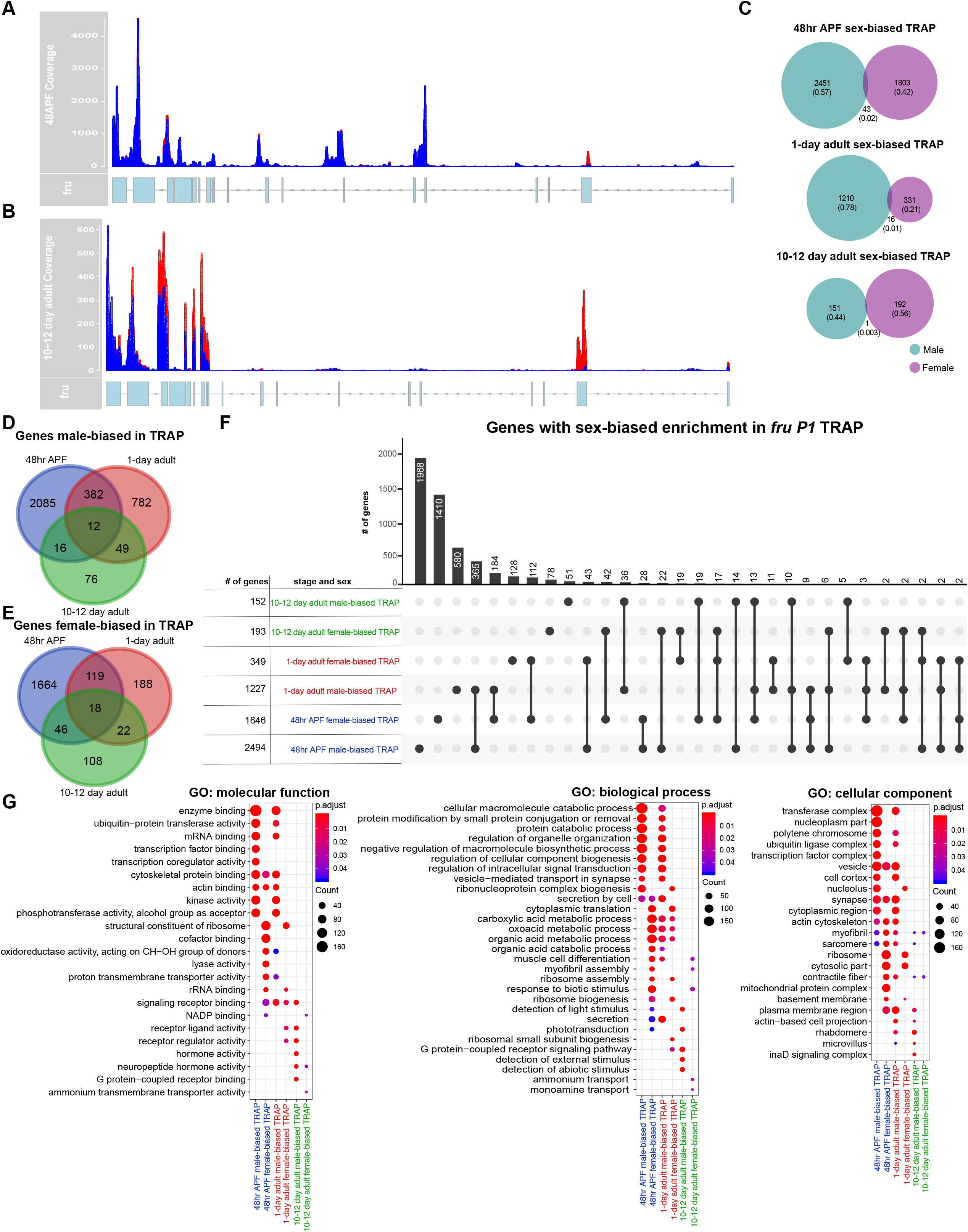
TRAP sequencing read coverage of *fruitless* in 48hr APF and 10-12 day data sets and analysis of sex-biased TRAP genes. **(A)** Average read coverage in 48hr APF TRAP and (**B**) 10-12 day adult TRAP for males (blue) and females (red) across the *fruitless* locus. The 5’ end of *fruitless* is located on the right and transcription runs in the direction of the arrows on the gene structure (right to left). **(C)** Sex-biased TRAP genes were identified by comparing expression between the male and female TRAP samples. For a gene to be considered sex-biased in the TRAP comparison, at least one exon needed to be more highly expressed in the TRAP sample of one sex (FDR<0.2). Comparison of male-biased TRAP genes (teal) and female-biased TRAP genes (purple) for all time points examined. For each Venn diagram category the number of genes and the proportion of the total genes (in parentheses) in each panel is shown. (**D-E**) Venn diagrams comparing sex-biased TRAP genes across time points within each sex. (**F**) Upset intersection plot of sex-biased TRAP genes from 48hr APF (blue), 1-day adult (red) (Newell et al., 2016) and 10-12 day adult (green) time points (Conway et al., 2017). n=4-5 biological replicates per condition. (**G**) Gene ontology (GO) enrichment analysis for sex-biased TRAP genes. The GO categories are molecular function, biological process, and cellular component. The GO terms shown in the plots are the top ten most significantly enriched terms for each list (non-redundant shown; Benjamini-Hochberg, p<0.05). The size of each dot indicates the number of genes (count) and the color indicates the p value (p.adjust). The GO information is in **Supplemental Table 4** and gene lists are in **Supplemental Table 5**.

**Supplemental Figure 6.**
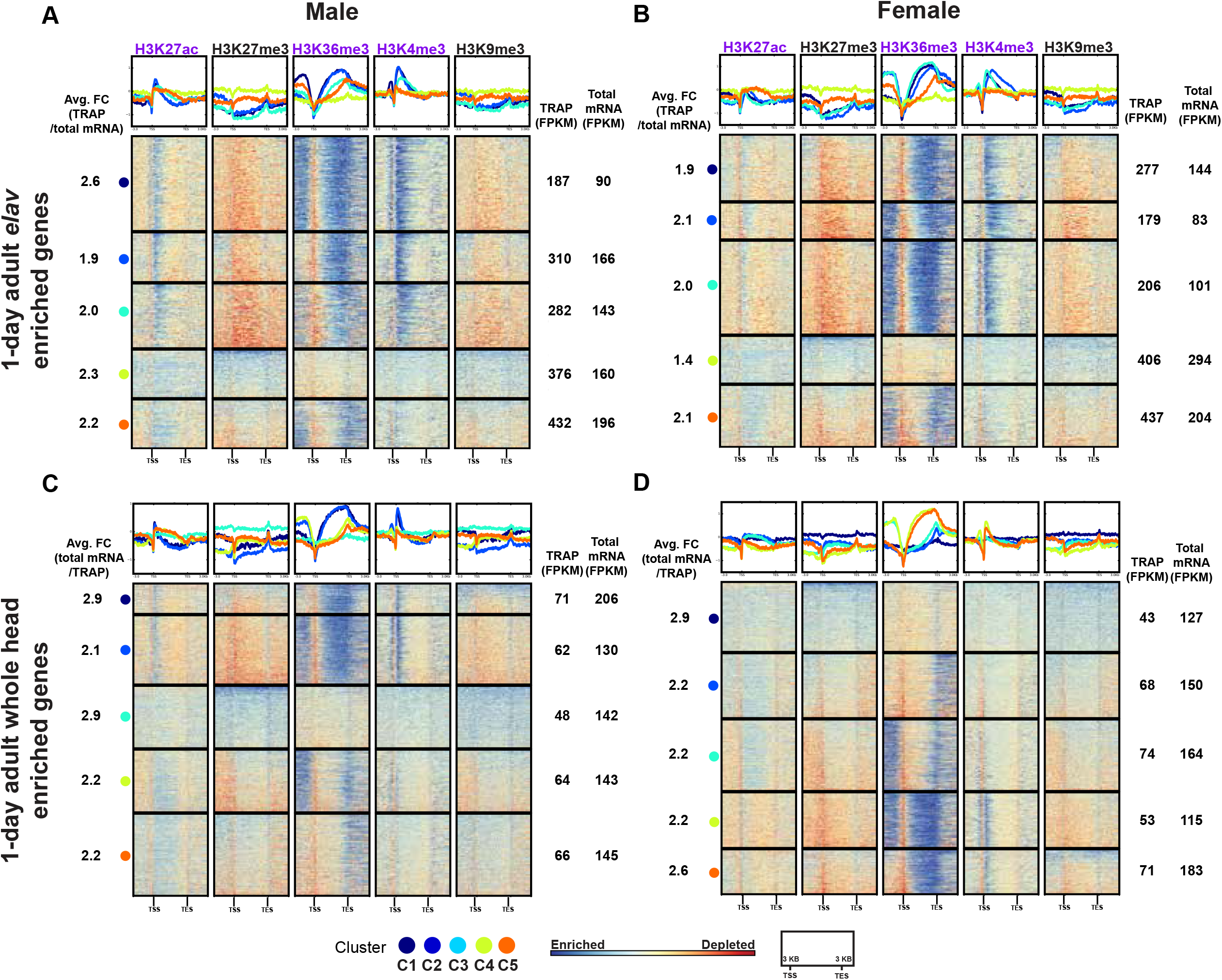
Hierarchical clustering of histone modification distributions for elav TRAP-enriched genes. For detailed description of heatmaps see **Figure 3**. Heatmaps for *elav* chromatin data from 1-day adults based on *elav* TRAP data enrichments (Thomas et al., 2012). **(A-B)** Heatmap for genes that are TRAP-enriched in 1-day adult *elav* neurons. **(C-D)** Heatmap for genes enriched in total mRNA. For each cluster of genes, the average fold-enrichment (Avg FC) of gene expression, indicated on the left (TRAP-enrichment), was calculated. For each cluster of genes, the average expression is indicated on the right for the TRAP and total mRNA data. Average fold-enrichment in gene expression and average expression in fragments per kilobase per million (FPKM) was calculated per cluster. The average was from expression data for genes that were significantly enriched in *elav* neurons (TRAP) from both sexes, or whole head tissue (total mRNA), averaged over both sexes (FDR<0.2; Thomas et al., 2012). Gene lists for each cluster are provided in **Supplemental Table 2**.

**Supplemental Figure 7.**
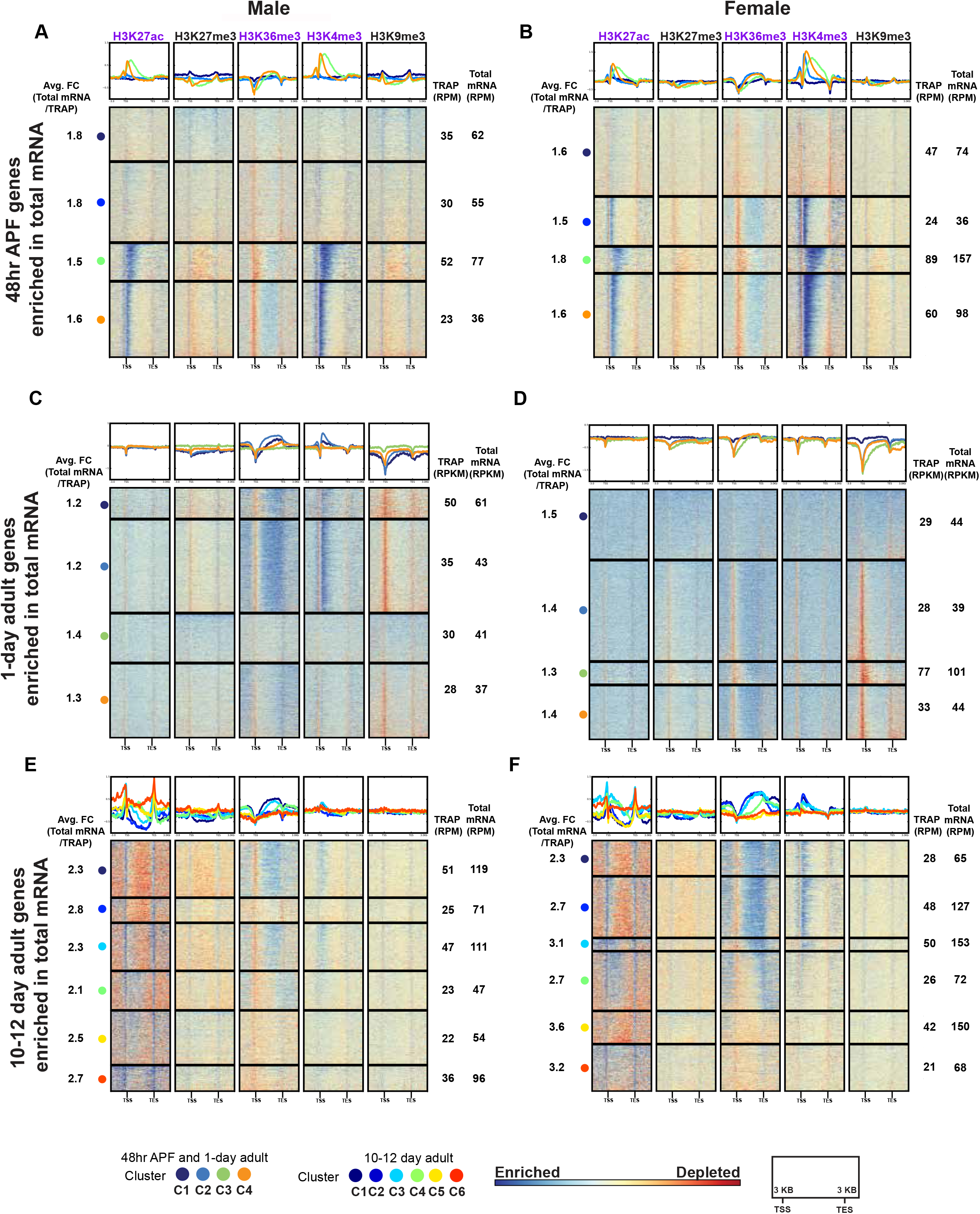
Hierarchical clustering of histone modification distributions for total mRNA-enriched genes. Hierarchal clustering reveals shared combinations of histone modifications for total mRNA-enriched genes relative to TRAP, generated using deepTools. For detailed description of heatmaps see **Figure 3**. Heatmaps for *fru P1* chromatin data from 48hr APF **(A-B)**, 1-day adults **(C-D)**, and 10-12 day adults **(E-F)**. Gene lists for each cluster are provided in **Supplemental Table 2**. For each cluster of genes, the average fold-enrichment (Avg FC) of gene expression is indicated on the left (total mRNA-enrichment), calculated at the exon level. For each cluster of genes, the average expression is indicated on the right for the TRAP and total mRNA data, calculated at the exon level. The 48hr APF and 10-12 day adult gene expression levels are provided as reads per million (RPM). The 1-day adult time point gene expression levels are provided as reads per kilobase per million (RPKM) (Newell et al., 2016). For TRAP and total mRNA expression data see **Supplemental Table 5**.

**Supplemental Figure 8.**
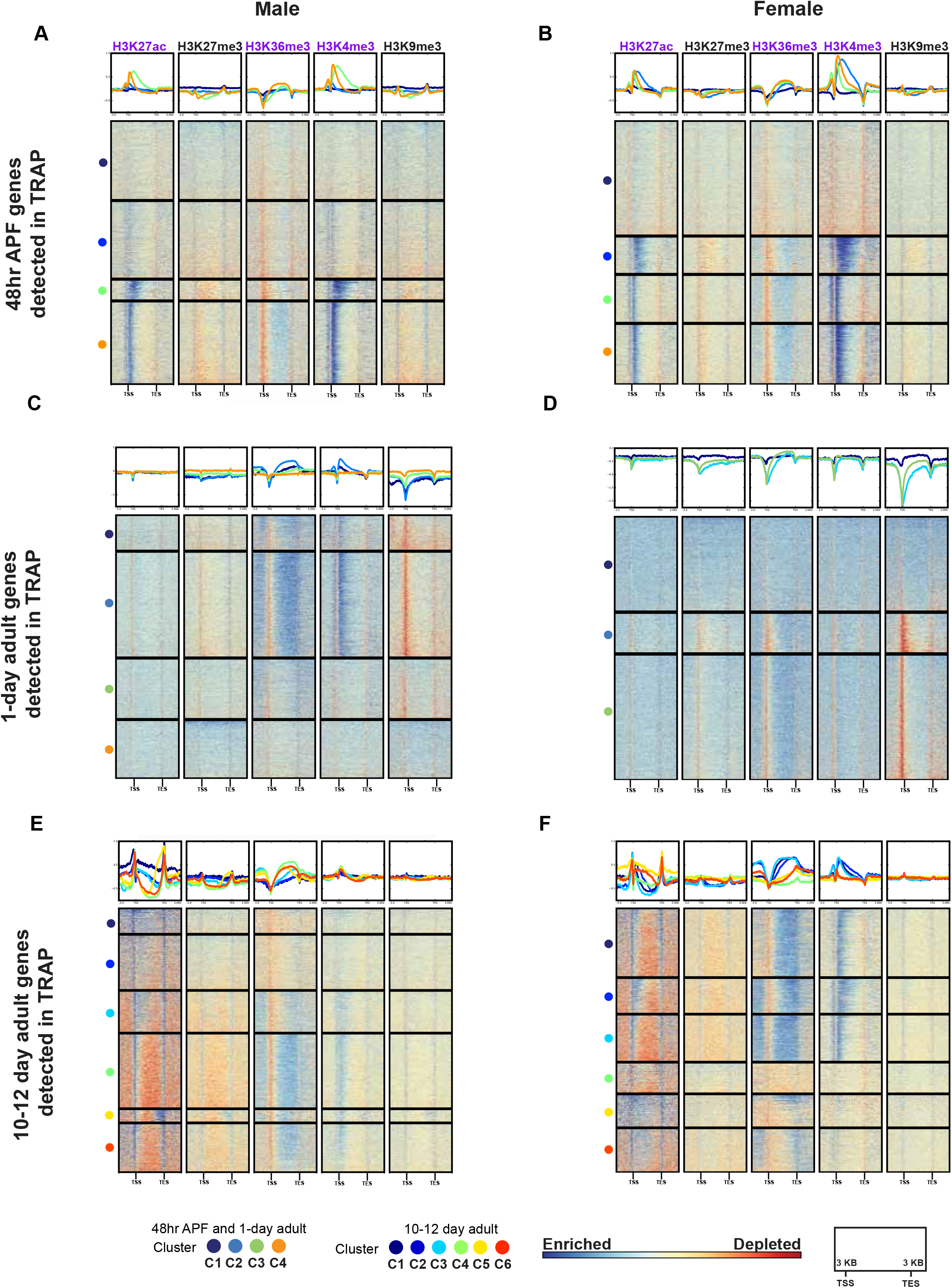
Hierarchical clustering of histone modification distributions for TRAP-detected genes. Hierarchal clustering reveals shared combinations of histone modifications for genes detected in expression data sets, generated using deepTools. For detailed description of heatmaps see **Figure 3**. Heatmaps for *fru P1* chromatin data from 48hr APF **(A-B)**, 1-day adults **(C-D)**, and 10-12 day adults **(E-F)**. A gene was considered detected in our 48hr APF or 10-12 day adult TRAP libraries if one or more exons were detected based on our initial filtering criteria (see methods). Genes detected in 1-day adult TRAP were those identified in Newell et al., 2016. Gene lists for each cluster are provided in **Supplemental Table 2**.

**Supplemental Figure 9.**
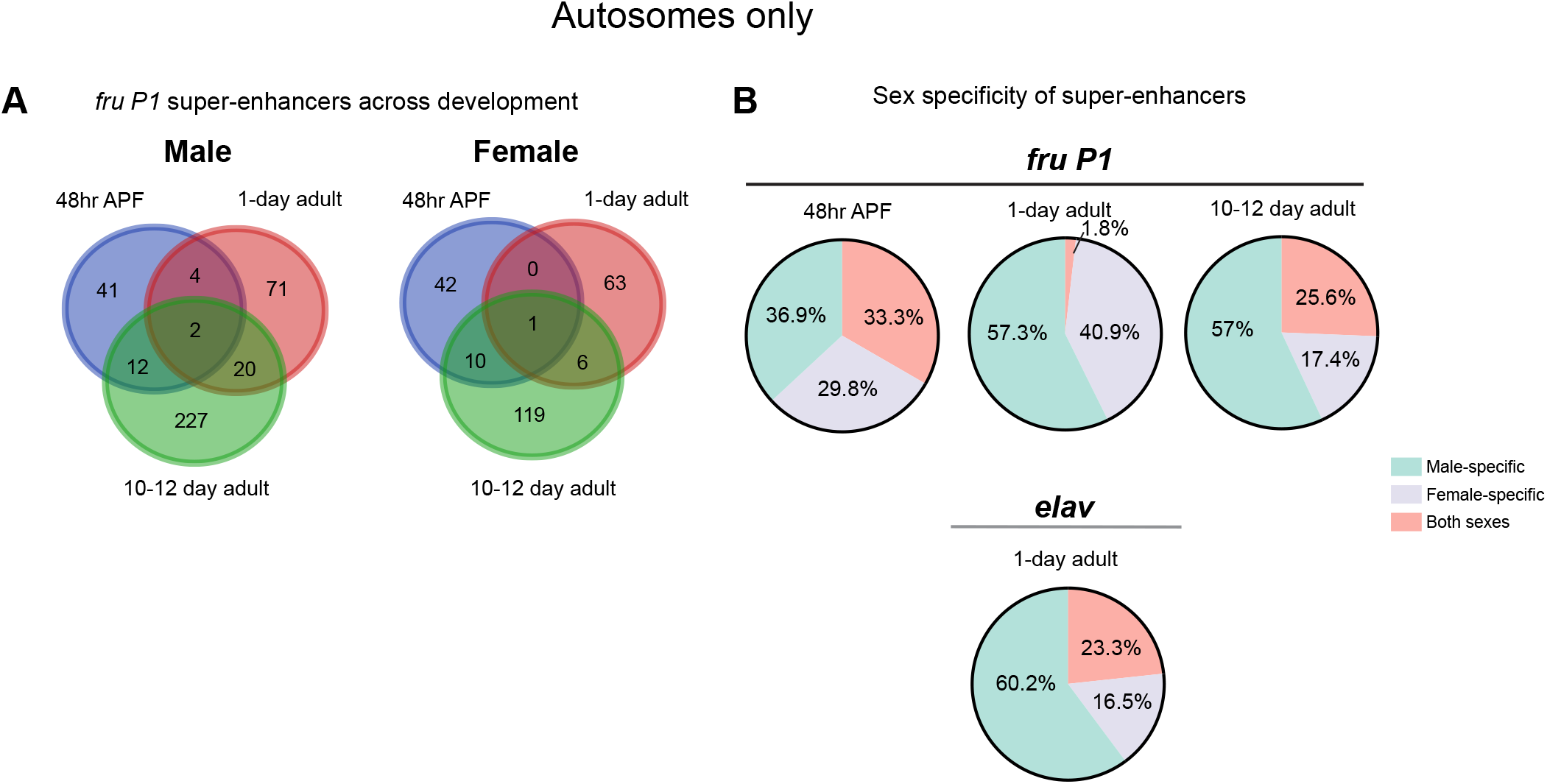
Super-enhancers identified based on H3K27ac peaks in *fru P1*- and *elav* neuron chromatin data sets for autosomes. **(A)** Venn diagrams showing the overlap between *fru P1* SE-containing genes identified on autosomes across time points for males (left) and females (right). **(B)** Percentages of SE-containing genes for each stage and neuron type (*fru P1* or *elav*) that are male-specific, female-specific or occur in both sexes.

## References

1. Dulac C. Brain function and chromatin plasticity. Nature. 2010;465(7299):728–35.

2. Graff J, Tsai LH. Histone acetylation: molecular mnemonics on the chromatin. Nature Reviews Neuroscience. 2013;14(2):97–111.

3. Crepaldi L, Riccio A. Chromatin learns to behave. Epigenetics. 2009;4(1):23–6.

4. Opachaloemphan C, Yan H, Leibholz A, Desplan C, Reinberg D. Recent Advances in Behavioral (Epi) Genetics in Eusocial Insects. Annual Review of Genetics, Vol 52. 2018;52:489–510.

5. Anreiter I, Biergans SD, Sokolowski MB. Epigenetic regulation of behavior in Drosophila melanogaster. Current Opinion in Behavioral Sciences. 2019;25:44–50.

6. Yamamoto D, Sato K, Koganezawa M. Neuroethology of male courtship in Drosophila: from the gene to behavior. Journal of Comparative Physiology a-Neuroethology Sensory Neural and Behavioral Physiology. 2014;200(4):251–64.

7. Dauwalder B. THE ROLES OF FRUITLESS AND DOUBLESEX IN THE CONTROL OF MALE COURTSHIP. Recent Advances in the Use of Drosophila in Neurobiology and Neurodegeneration. 2011;99:87–105.

8. Kubli E, Bopp D. Sexual Behavior: How Sex Peptide Flips the Postmating Switch of Female Flies. Current Biology. 2012;22(13):R520–R2.

9. Laturney M, Billeter JC. Neurogenetics of Female Reproductive Behaviors in Drosophila melanogaster. Advances in Genetics, Vol 85. 2014;85:1–108.

10. Burtis KC, Baker BS. DROSOPHILA DOUBLESEX GENE CONTROLS SOMATIC SEXUAL-DIFFERENTIATION BY PRODUCING ALTERNATIVELY SPLICED MESSENGER-RNAS ENCODING RELATED SEX-SPECIFIC POLYPEPTIDES. Cell. 1989;56(6):997–1010.

11. Ryner LC, Goodwin SF, Castrillon DH, Anand A, Villella A, Baker BS, et al. Control of male sexual behavior and sexual orientation in Drosophila by the fruitless gene. Cell. 1996;87(6):1079–89.

12. Ito H, Fujitani K, Usui K, ShimizuNishikawa K, Tanaka S, Yamamoto D. Sexual orientation in Drosophila is altered by the satori mutation in the sex-determination gene fruitless that encodes a zinc finger protein with a BTB domain. P Natl Acad Sci USA. 1996;93(18):9687–92.

13. Villella A, Hall JC. Courtship anomalies caused by doublesex mutations in Drosophila melanogaster. Genetics. 1996;143(1):331–44.

14. Robinett CC, Vaughan AG, Knapp JM, Baker BS. Sex and the Single Cell. II. There Is a Time and Place for Sex. Plos Biology. 2010;8(5).

15. Rideout EJ, Dornan AJ, Neville MC, Eadie S, Goodwin SF. Control of sexual differentiation and behavior by the doublesex gene in Drosophila melanogaster. Nature Neuroscience. 2010;13(4):458–U79.

16. Lee G, Hall JC, Park JH. Doublesex gene expression in the central nervous system of Drosophila melanogaster. Journal of Neurogenetics. 2002;16(4):229–48.

17. Lee G, Foss M, Goodwin SF, Carlo T, Taylor BJ, Hall JC. Spatial, temporal, and sexually dimorphic expression patterns of the fruitless gene in the Drosophila central nervous system. Journal of Neurobiology. 2000;43(4):404–26.

18. Sanders LE, Arbeitman MN. Doublesex establishes sexual dimorphism in the Drosophila central nervous system in an isoform-dependent manner by directing cell number. Developmental Biology. 2008;320(2):378–90.

19. Manoli DS, Foss M, Villella A, Taylor BJ, Hall JC, Baker BS. Male-specific fruitless specifies the neural substrates of Drosophila courtship behaviour. Nature. 2005;436(7049):395–400.

20. Stockinger P, Kvitsiani D, Rotkopf S, Tirian L, Dickson BJ. Neural circuitry that governs Drosophila male courtship behavior. Cell. 2005;121(5):795–807.

21. Hall JC. COURTSHIP AMONG MALES DUE TO A MALE-STERILE MUTATION IN DROSOPHILA-MELANOGASTER. Behavior Genetics. 1978;8(2):125–41.

22. Anand A, Villella A, Ryner LC, Carlo T, Goodwin SF, Song HJ, et al. Molecular genetic dissection of the sex-specific and vital functions of the Drosophila melanogaster sex determination gene fruitless. Genetics. 2001;158(4):1569–95.

23. Goodwin SF, Taylor BJ, Villella A, Foss M, Ryner LC, Baker BS, et al. Aberrant splicing and altered spatial expression patterns in fruitless mutants of Drosophila melanogaster. Genetics. 2000;154(2):725–45.

24. Villella A, Gailey DA, Berwald B, Ohshima S, Barnes PT, Hall JC. Extended reproductive roles of the fruitless gene in Drosophila melanogaster revealed by behavioral analysis of new fru mutants. Genetics. 1997;147(3):1107–30.

25. Demir E, Dickson BJ. fruitless splicing specifies male courtship behavior in Drosophila. Cell. 2005;121(5):785–94.

26. Kvitsiani D, Dickson BJ. Shared neural circuitry for female and male sexual behaviours in Drosophila. Current Biology. 2006;16(10):R355–R6.

27. Kimura KI, Ote M, Tazawa T, Yamamoto D. Fruitless specifies sexually dimorphic neural circuitry in the Drosophila brain. Nature. 2005;438(7065):229–33.

28. Belote JM, Baker BS. SEXUAL-BEHAVIOR - ITS GENETIC-CONTROL DURING DEVELOPMENT AND ADULTHOOD IN DROSOPHILA-MELANOGASTER. Proceedings of the National Academy of Sciences of the United States of America. 1987;84(22):8026–30.

29. Arthur BI, Jallon JM, Caflisch B, Choffat Y, Nothiger R. Sexual behaviour in Drosophila is irreversibly programmed during a critical period. Current Biology. 1998;8(21):1187–90.

30. Hueston CE, Olsen D, Li QY, Okuwa S, Peng B, Wu JN, et al. Chromatin Modulatory Proteins and Olfactory Receptor Signaling in the Refinement and Maintenance of Fruitless Expression in Olfactory Receptor Neurons. Plos Biology. 2016;14(4).

31. Zhao SH, Deanhardt B, Barlow GT, Schleske PG, Rossi AM, Volka PC. Chromatin-based reprogramming of a courtship regulator by concurrent pheromone perception and hormone signaling. Science Advances. 2020;6(21).

32. Sethi S, Lin HH, Shepherd AK, Volkan PC, Su CY, Wang JW. Social Context Enhances Hormonal Modulation of Pheromone Detection in Drosophila. Current Biology. 2019;29(22):3887-+.

33. Ito H, Sato K, Koganezawa M, Ote M, Matsumoto K, Hama C, et al. Fruitless Recruits Two Antagonistic Chromatin Factors to Establish Single-Neuron Sexual Dimorphism. Cell. 2012;149(6):1327–38.

34. Handley A, Schauer T, Ladurner AG, Margulies CE. Designing Cell-Type-Specific Genome-wide Experiments. Molecular Cell. 2015;58(4):621–31.

35. van den Ameelel J, Krautz R, Brand AH. TaDa! Analysing cell type-specific chromatin in vivo with Targeted DamID. Current Opinion in Neurobiology. 2019;56:160–6.

36. Bannister AJ, Kouzarides T. Regulation of chromatin by histone modifications. Cell Research. 2011;21(3):381–95.

37. Cao R, Wang LJ, Wang HB, Xia L, Erdjument-Bromage H, Tempst P, et al. Role of histone H3 lysine 27 methylation in polycomb-group silencing. Science. 2002;298(5595):1039–43.

38. Creyghton MP, Cheng AW, Welstead GG, Kooistra T, Carey BW, Steine EJ, et al. Histone H3K27ac separates active from poised enhancers and predicts developmental state. Proceedings of the National Academy of Sciences of the United States of America. 2010;107(50):21931–6.

39. Fischle W, Wang YM, Jacobs SA, Kim YC, Allis CD, Khorasanizadeh S. Molecular basis for the discrimination of repressive methyl-lysine marks in histone H3 bv Polvcomb and HP1 chromodomains. Genes & Development. 2003;17(15):1870–81.

40. Kolasinska-Zwierz P, Down T, Latorre I, Liu T, Liu XS, Ahringer J. Differential chromatin marking of introns and expressed exons by H3K36me3. Nature Genetics. 2009;41(3):376–81.

41. Thomas A, Lee PJ, Dalton JE, Nomie KJ, Stoica L, Costa-Mattioli M, et al. A Versatile Method for Cell-Specific Profiling of Translated mRNAs in Drosophila. Plos One. 2012;7(7).

42. Newell NR, New FN, Dalton JE, McIntyre LM, Arbeitman MN. Neurons That Underlie Drosophila melanogaster Reproductive Behaviors: Detection of a Large Male-Bias in Gene Expression in fruitless-Expressing Neurons. G3-Genes Genomes Genetics. 2016;6(8):2455–65.

43. Brand AH, Perrimon N. TARGETED GENE-EXPRESSION AS A MEANS OF ALTERING CELL FATES AND GENERATING DOMINANT PHENOTYPES. Development. 1993;118(2):401–15.

44. Duffy JB. GAL4 system in Drosophila: A fly geneticist's Swiss army knife. Genesis. 2002;34(1-2):1–15.

45. Beckett D, Kovaleva E, Schatz PJ. A minimal peptide substrate in biotin holoenzyme synthetase-catalyzed biotinylation. Protein Science. 1999;8(4):921–9.

46. Riddle NC, Elgin SCR. The Drosophila Dot Chromosome: Where Genes Flourish Amidst Repeats. Genetics. 2018;210(3):757–72.

47. Treangen TJ, Salzberg SL. Repetitive DNA and next-generation sequencing: computational challenges and solutions. Nature Reviews Genetics. 2012;13(1):36–46.

48. Hall JC. THE MATING OF A FLY. Science. 1994;264(5166):1702–14.

49. Hing ALY, Carlson JR. Male-male courtship behavior induced by ectopic expression of the Drosophila white gene: Role of sensory function and age. Journal of Neurobiology. 1996;30(4):454–64.

50. Yamamoto D, Jallon JM, Komatsu A. Genetic dissection of sexual behavior in Drosophila melanogaster. Annual Review of Entomology. 1997;42:551–85.

51. Zhang SD, Odenwald WF. MISEXPRESSION OF THE WHITE (w) GENE TRIGGERS MALE-MADE COURTSHIP IN DROSOPHILA. Proceedings of the National Academy of Sciences of the United States of America. 1995;92(12):5525–9.

52. Ramirez F, Ryan DP, Gruning B, Bhardwaj V, Kilpert F, Richter AS, et al. deepTools2: a next generation web server for deep-sequencing data analysis. Nucleic Acids Research. 2016;44(W1):W160–W5.

53. Kharchenko PV, Alekseyenko AA, Schwartz YB, Minoda A, Riddle NC, Ernst J, et al. Comprehensive analysis of the chromatin landscape in Drosophila melanogaster. Nature. 2011;471(7339):480-+.

54. Kouzarides T. Chromatin modifications and their function. Cell. 2007;128(4):693–705.

55. Zhang Y, Liu T, Meyer CA, Eeckhoute J, Johnson DS, Bernstein BE, et al. Model-based Analysis of ChIP-Seq (MACS). Genome Biology. 2008;9(9).

56. Pal K, Forcato M, Jost D, Sexton T, Vaillant C, Salviato E, et al. Global chromatin conformation differences in the Drosophila dosage compensated chromosome X. Nature Communications. 2019;10.

57. Gelbart ME, Larschan E, Peng SY, Park PJ, Kuroda MI. Drosophila MSL complex globally acetylates H4K16 on the male X chromosome for dosage compensation. Nature Structural & Molecular Biology. 2009;16(8):825–U47.

58. Negre N, Brown CD, Ma LJ, Bristow CA, Miller SW, Wagner U, et al. A cis-regulatory map of the Drosophila genome. Nature. 2011;471(7339):527–31.

59. Marsano RM, Giordano E, Messina G, Dimitri P. A New Portrait of Constitutive Heterochromatin: Lessons from Drosophila melanogaster. Trends in Genetics. 2019;35(9):615–31.

60. Lyne R, Smith R, Rutherford K, Wakeling M, Varley A, Guillier F, et al. FlyMine: an integrated database for Drosophila and Anopheles genomics. Genome Biology. 2007;8(7).

61. Shen LS M., GeneOverlap: Test and visualize gene overlaps. 2019.

62. Dalton JE, Fear JM, Knott S, Baker BS, McIntyre LM, Arbeitman MN. Male-specific Fruitless isoforms have different regulatory roles conferred by distinct zinc finger DNA binding domains. Bmc Genomics. 2013;14.

63. Neville MC, Nojima T, Ashley E, Parker DJ, Walker J, Southall T, et al. Male-Specific Fruitless Isoforms Target Neurodevelopmental Genes to Specify a Sexually Dimorphic Nervous System. Current Biology. 2014;24(3):229–41.

64. Chen J, Jin S, Cao J, Peng Q, Pan Y. <em>fruitless</em> tunes functional flexibility of courtship circuitry during development. bioRxiv. 2020:2020.06.20.163055.

65. Conway JR, Lex A, Gehlenborg N. UpSetR: an R package for the visualization of intersecting sets and their properties. Bioinformatics. 2017;33(18):2938–40.

66. Brovero SG, Fortier JC, Hu H, Lovejoy PC, Newell NR, Palmateer CM, et al. Neurogenetic and genomic approaches reveal roles for Dpr/DIP cell adhesion molecules in <em>Drosophila</em> reproductive behavior. bioRxiv. 2020:2020.10.02.323477.

67. Penalva LOF, Sanchez L. RNA binding protein sex-lethal (Sxl) and control of Drosophila sex determination and dosage compensation. Microbiology and Molecular Biology Reviews. 2003;67(3):343-+.

68. Whyte WA, Orlando DA, Hnisz D, Abraham BJ, Lin CY, Kagey MH, et al. Master Transcription Factors and Mediator Establish Super-Enhancers at Key Cell Identity Genes. Cell. 2013;153(2):307–19.

69. Loven J, Hoke HA, Lin CY, Lau A, Orlando DA, Vakoc CR, et al. Selective Inhibition of Tumor Oncogenes by Disruption of Super-Enhancers. Cell. 2013;153(2):320–34.

70. Yu JY, Kanai MI, Demir E, Jefferis G, Dickson BJ. Cellular Organization of the Neural Circuit that Drives Drosophila Courtship Behavior. Current Biology. 2010;20(18):1602–14.

71. Gohl DM, Silies MA, Gao XJJ, Bhalerao S, Luongo FJ, Lin CC, et al. A versatile in vivo system for directed dissection of gene expression patterns. Nature Methods. 2011;8(3):231–U71.

72. Diao FQ, Ironfield H, Luan HJ, Diao FC, Shropshire WC, Ewer J, et al. Plug-and-Play Genetic Access to Drosophila Cell Types using Exchangeable Exon Cassettes. Cell Reports. 2015;10(8):1410–21.

73. Goldman TD, Arbeitman MN. Genomic and functional studies of Drosophila sex hierarchy regulated gene expression in adult head and nervous system tissues. Plos Genetics. 2007;3(11):2278–95.

74. Vernes SC. Genome wide identification of Fruitless targets suggests a role in upregulating genes important for neural circuit formation. Scientific Reports. 2014;4.

75. Arbeitman MN, Fleming AA, Siegal ML, Null BH, Baker BS. A genomic analysis of Drosophila somatic sexual differentiation and its regulation. Development. 2004;131(9):2007–21.

76. Kwasnieski JC, Fiore C, Chaudhari HG, Cohen BA. High-throughput functional testing of ENCODE segmentation predictions. Genome Research. 2014;24(10):1595–602.

77. Arvey A, Agius P, Noble WS, Leslie C. Sequence and chromatin determinants of cell-type-specific transcription factor binding. Genome Research. 2012;22(9):1723–34.

78. Brown EJ, Bachtrog D. The chromatin landscape of Drosophila: comparisons between species, sexes, and chromosomes. Genome Research. 2014;24(7):1125–37.

79. Pulecio J, Verma N, Mejia-Ramirez E, Huangfu D, Raya A. CRISPR/Cas9-Based Engineering of the Epigenome. Cell Stem Cell. 2017;21(4):431–47.

80. Brezgin S, Kostyusheva A, Kostyushev D, Chulanov V. Dead Cas Systems: Types, Principles, and Applications. International Journal of Molecular Sciences. 2019;20(23).

81. Griffith LC, Ejima A. Courtship learning in Drosophila melanogaster: Diverse plasticity of a reproductive behavior. Learning & Memory. 2009;16(12):743–50.

82. Andrew DJ, Chen EH, Manoli DS, Ryner LC, Arbeitman MN. Sex and the Single Fly: A Perspective on the Career of Bruce S. Baker. Genetics. 2019;212(2):365–76.

83. Bolger AM, Lohse M, Usadel B. Trimmomatic: a flexible trimmer for Illumina sequence data. Bioinformatics. 2014;30(15):2114–20.

84. dos Santos G, Schroeder AJ, Goodman JL, Strelets VB, Crosby MA, Thurmond J, et al. FlyBase: introduction of the Drosophila melanogaster Release 6 reference genome assembly and large-scale migration of genome annotations. Nucleic Acids Research. 2015;43(D1):D690–D7.

85. Trapnell C, Pachter L, Salzberg SL. TopHat: discovering splice junctions with RNA-Seq. Bioinformatics. 2009;25(9):1105–11.

86. Li H, Handsaker B, Wysoker A, Fennell T, Ruan J, Homer N, et al. The Sequence Alignment/Map format and SAMtools. Bioinformatics. 2009;25(16):2078–9.

87. Dobin A, Davis CA, Schlesinger F, Drenkow J, Zaleski C, Jha S, et al. STAR: ultrafast universal RNA-seq aligner. Bioinformatics. 2013;29(1):15–21.

88. Kellis M, Wold B, Snyder MP, Bernstein BE, Kundaje A, Marinov GK, et al. Defining functional DNA elements in the human genome. Proceedings of the National Academy of Sciences of the United States of America. 2014;111(17):6131–8.

89. Ross-Innes CS, Stark R, Teschendorff AE, Holmes KA, Ali HR, Dunning MJ, et al. Differential oestrogen receptor binding is associated with clinical outcome in breast cancer. Nature. 2012;481(7381):389–U177.

90. Yu GC, Wang LG, He QY. ChIPseeker: an R/Bioconductor package for ChIP peak annotation, comparison and visualization. Bioinformatics. 2015;31(14):2382–3.

91. Liao Y, Smyth GK, Shi W. featureCounts: an efficient general purpose program for assigning sequence reads to genomic features. Bioinformatics. 2014;30(7):923–30.

92. Robinson MD, McCarthy DJ, Smyth GK. edgeR: a Bioconductor package for differential expression analysis of digital gene expression data. Bioinformatics. 2010;26(1):139–40.

93. Yu GC, Wang LG, Han YY, He QY. clusterProfiler: an R Package for Comparing Biological Themes Among Gene Clusters. Omics-a Journal of Integrative Biology. 2012;16(5):284–7.

94. Cichewicz K, Hirsh J. ShinyR-DAM: a program analyzing Drosophila activity, sleep and circadian rhythms. Communications Biology. 2018;1.

95. Hulsen T, de Vlieg J, Alkema W. BioVenn - a web application for the comparison and visualization of biological lists using area-proportional Venn diagrams. Bmc Genomics. 2008;9.

